# Osmo-hydraulic volume regulation ensures robust endothelial-to-haematopoietic transition

**DOI:** 10.64898/2026.04.09.711986

**Authors:** Igor Kondrychyn, Yan Chen, Rajrishi Kumar, Jason da Silva, Haymar Wint, Guihua Chen, Koichi Kawakami, Eoin McEvoy, Li-Kun Phng

## Abstract

Haematopoietic stem and progenitor cells (HSPCs) emerge from specialised haemogenic endothelial cells (HECs) in the ventral wall of the dorsal aorta (VDA) through a process known as endothelial-to-haematopoietic transition (EHT). During EHT, elongated HECs undergo actomyosin-driven rounding and extrusion, a mechanically demanding transformation that requires cell integrity to be maintained. Here, we identify an osmo-hydraulic volume-regulation mechanism that enables HECs to adapt to and withstand this mechanical challenge. Haemodynamic forces increase Piezo-dependent Ca²⁺ activity in ventral endothelial cells, while Piezo activation promotes HEC swelling. Combined genetic and pharmacological analyses support a pathway in which Piezo-mediated swelling triggers VRAC-dependent osmolyte efflux and Aqp1a.1-mediated water efflux to reduce HEC volume. Disrupting either osmolyte or water efflux causes excessive HEC swelling, compromises HEC integrity and reduces HSPC production. Together, our findings establish osmo-hydraulic volume regulation as a mechanism that preserves cellular robustness during EHT and support a model in which aquaporins act as pressure-relief valves, dissipating intracellular hydrostatic pressure to sustain HEC survival and definitive haematopoiesis.

## Introduction

The dorsal aorta (DA) is the first artery to form during embryogenesis. Beyond its essential role in distributing blood from the heart to the embryonic trunk, the DA serves as a major developmental source of endothelial cells (ECs) for trunk vascular expansion and of multipotent haematopoietic stem and progenitor cells (HSPCs), which possess the capacity for self-renewal and multilineage differentiation of the blood system^1,2^. Spatial patterning within the DA underlies these distinct outputs. At the dorsal roof, a subset of ECs becomes specified as tip cells that emerge to form intersegmental vessels (ISVs) ^3,4^. ISVs subsequently elongate through the dorsal migration of tip cells, the addition of endothelial stalk cells originating from the DA, and cell proliferation ^4–6^. In contrast, at the ventral floor of the DA (VDA), specialised haemogenic endothelial cells (HECs) generate HSPCs that extrude either as clusters into the aortic lumen in amniotes ^7,8^, or as individual cells into the subaortic space in the zebrafish ^7,9,10^. This process, termed endothelial-to-haematopoietic transition (EHT), occurs within a narrow developmental window and is governed by coordinated transcriptional and biophysical cues.

The zebrafish has been vital in elucidating the stepwise mechanisms underlying EHT. Between ∼ 12 and 16 hpf, arterial-fated angioblasts migrate from the lateral plate mesoderm (LPM) to the embryonic midline, where they assemble to form the DA^11^. Subsequently, a subpopulation of ECs acquires haemogenic identity through a transcriptional switch characterised by the downregulating endothelial genes such as *kdr* and *cdh5* and the upregulation of the haematopoietic transcription factors *gata2b*, *runx1* and c*myb* to become HECs ^1,12^. Gata2b acts upstream of Runx1^13^, whose activity is indispensable for haematopoietic fate commitment and for the execution of EHT in both zebrafish and mice^14–17^. Several signalling pathways contribute to HEC specification, including Notch and Wnt signalling ^18,19^, while Apelin signalling acts to maintain arterial identity, such that its suppression enhances EC-to-HEC conversion^20^. In addition to transcriptional regulation, EHT is influenced by the mechanical environment of the DA^12^. Notably, following the onset of cardiac contraction, haemodynamic forces such as shear stress and cyclic stretch promote EHT and HSPC generation by inducing *runx1* expression^21,22^. Cyclic stretching of the endothelium further activates YAP signalling, which also induces *runx1* expression to sustain the production of HSPCs ^23^.

A defining feature of EHT is the extensive morphological transition that HECs undergo. Live imaging in the zebrafish has revealed the relocation of HECs from the dorsal roof to the ventral floor of the DA, where they adopt an elongated morphology before rounding up, undergoing apical invagination and extruding from the vessel wall into the subaortic space between ∼ 28 and 48 hpf ^24,25^. This transition is driven by actomyosin contractility, establishment of apicobasal polarity and junction remodelling between HECs and neighbouring arterial ECs^24,26^. In amniotes, EHT is further facilitated by intracellular vacuole formation^27^. The loss of cells from the VDA is subsequently compensated by the ventral movement and proliferation of ECs from the dorsal DA ^28^, thus maintaining vascular integrity during EHT.

Aquaporins are membrane-embedded channels that facilitate bidirectional water movement across cellular membranes, with the direction of flow determined by the osmotic gradient^29^. Emerging evidence indicates that aquaporins have roles beyond water homeostasis by regulating cell volume and shape to drive dynamic cell behaviours including migration ^30–34^ and tissue morphogenesis ^30,35^. We previously demonstrated that during zebrafish ISV development, endothelial aquaporins mediate water influx into tip cells, increasing cell volume, elevating local cytosolic hydrostatic pressure and promoting actin polymerisation to generate membrane protrusions that enhance tip cell emergence and migration from the DA ^30^. Given that HSPCs also arise from the DA and that EHT involves pronounced changes in cell shape and mechanics, we investigated whether aquaporins contribute to the regulation of HEC behaviour during EHT.

## Results

### Aqp1a.1 promotes definitive haematopoiesis

To examine the function of aquaporin in EHT, we utilised zebrafish lacking *aqp1a.1*, an aquaporin isoform that is highly expressed in arterial ECs of the DA ^30,36,37^. Analysis of *aqp1a.1^rk28/rk28^*;*Tg(fli1:H2B-EGFP);Tg(fli1:Lifeact-mCherry)* zebrafish at two stages (29 – 32 hpf and 48 – 52 hpf) shows a reduction in the number of rounded cells that have extruded from the VDA when compared to wild-type embryos (Supplementary Fig. 1a, b). Based on their rounded morphology, spatial location at the VDA and the timing of EHT, we inferred that these ECs represent HECs and nascent HSPCs, and that their number is reduced in the absence of Aqp1a.1. We verified this interpretation by employing a reporter for HECs and HSPCs, *Tg(gata2b:GAL4FF);Tg(UAS:EGFP)*, which recapitulates endogenous *gata2b* expression^38^, allowing direct visualisation of HECs in the DA. Quantification shows a significant decrease in *gata2b^+^* cells at the VDA in *aqp1a.1^rk28/rk28^* embryos when compared to wild-type and *aqp1a.1^+/rk28^* embryos at 28 – 31 (Fig. 1a, b) and 48 - 51 hpf (Fig. 1c, d). The decrease in HECs in *aqp1a.1^rk28/rk28^* embryos is further confirmed by a reduction in *runx1* (Fig. 1e, f) and *cmyb* (Fig. 1g, h) mRNA expression in the DA at 24 and 30 hpf, respectively.

**Figure 1.**
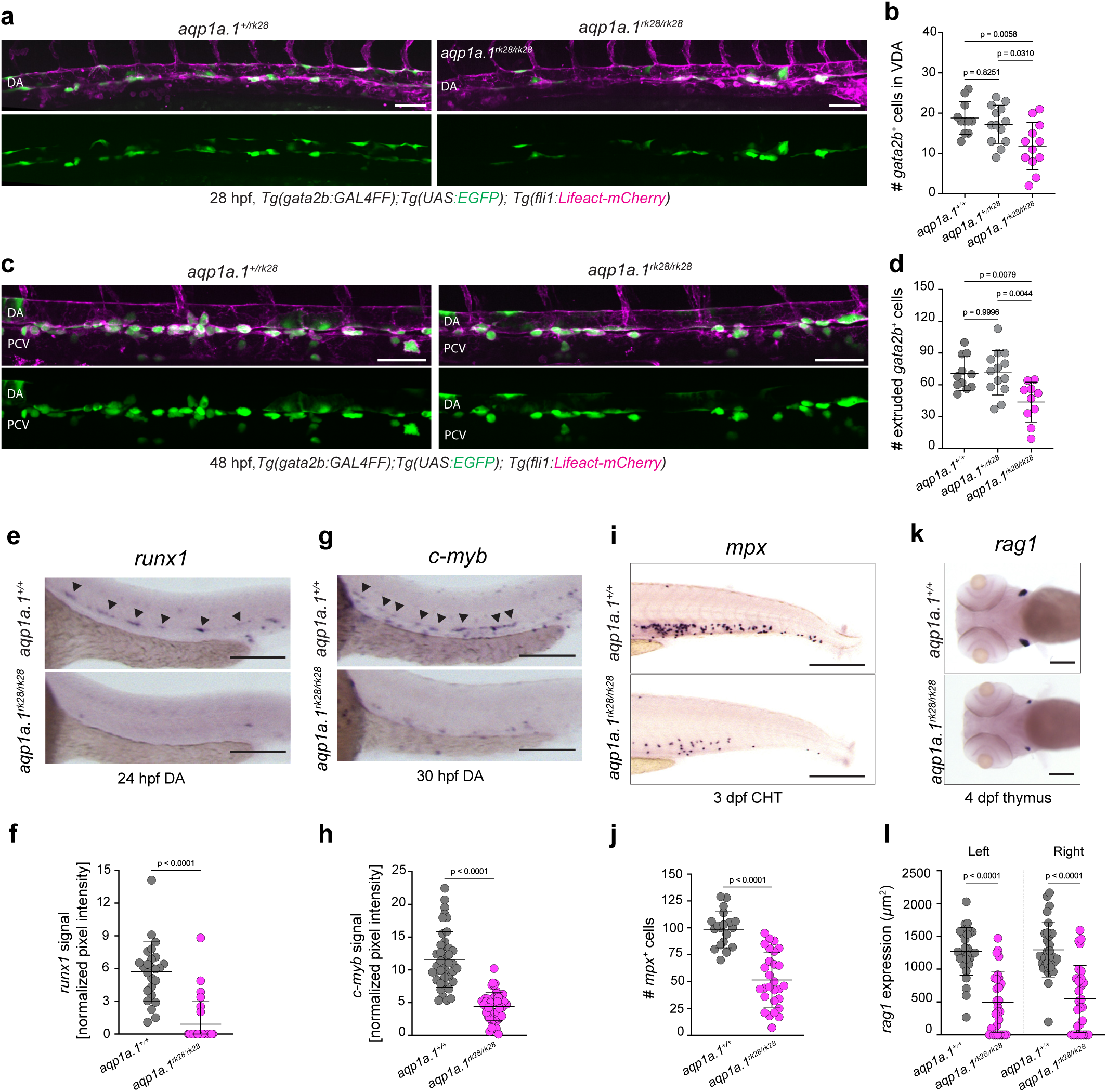
Aqp1a.1 promotes definitive haematopoiesis. **a** Representative maximum intensity projection images of *aqp1a.1^+/rk28^* and *aqp1a.1^rk28/rk28^* embryos in *Tg(gata2b:GAL4FF);Tg(UAS:EGFP);Tg(fli1:Lifeact-mCherry)* background at 28 hpf. **b** Quantification of *gata2b^+^* cells at VDA in *aqp1a.1^+/+^*, *aqp1a.1^+/rk28^* and *aqp1a.1^rk28/rk28^* embryos at 28-31 hpf (*aqp1a.1^+/+^*, n=11; *aqp1a.1^+/rk28^*, n=13; *aqp1a.1^rk28/rk28^*, n=12 embryos). **c** Representative maximum intensity projection images of *aqp1a.1^+/rk28^*and *aqp1a.1^rk28/rk28^* embryos in *Tg(gata2b:GAL4FF);Tg(UAS:EGFP);Tg(fli1:Lifeact-mCherry)* background at 48 hpf. **d** Quantification of extruded *gata2b^+^*cells in *aqp1a.1^+/+^*, *aqp1a.1^+/rk28^* and *aqp1a.1^rk28/rk28^*embryos at 48-51 hpf (*aqp1a.1^+/+^*, n=11; *aqp1a.1^+/rk28^*, n=13; *aqp1a.1^rk28/rk28^*, n=10 embryos). **e**, **g**, **i**, **k** Representative images of expression of *runx1* (**e**) and *c-myb* (**g**) in the DA (black arrowheads), *mpx* (**i**) in the CHT and *rag1* (**k**) in the thymus of *aqp1a.1^+/+^* and *aqp1a.1^rk28/rk28^* embryos at the indicated stages. **f**, **h**, **j**, **l** Quantification of *runx1* expression (**f**: *aqp1a.1^+/+^*, n=25; *aqp1a.1^rk28/rk28^*, n=28 embryos), *c-myb* expression (**h**: *aqp1a.1^+/+^*, n=32; *aqp1a.1^rk28/rk28^*, n=45 embryos), *mpx^+^* cell number (**j**: *aqp1a.1^+/+^*, n=20; *aqp1a.1^rk28/rk28^*, n=28 embryos) and *rag1* expression area (**l**: *aqp1a.1^+/+^*, n=29; *aqp1a.1^rk28/rk28^*, n=28 embryos) in *aqp1a.1^+/+^* and *aqp1a.1^rk28/rk28^* embryos. Data were collected from two independent experiments and presented as mean ± SD. Statistical significance was determined by ordinary one-way ANOVA with Sidak’s multiple comparisons test (**b**, **d** and **l**) and two-tailed unpaired *t*-test (**f**, **h**, **j**). CHT, caudal hematopoietic tissue; DA, dorsal aorta; PCV, posterior cardinal vein; VDA, ventral wall of the DA; hpf, hours post-fertilization. Scale bars, 50 μm (**a**, **c**), 150 μm (**e**, **g**, **k**) and 250 μm (**i**).

After extrusion from the VDA into the subaortic space, HSPCs enter the blood circulation through the posterior cardinal vein to colonise the caudal haematopoietic tissue (CHT), where they undergo clonal expansion and differentiation before migrating and seeding the thymus and kidney^39,40^. We therefore examined whether the decrease in HSPC number compromised the production of mature blood lineages. In *aqp1a.1^rk28/rk28^* zebrafish, there is a reduction in *mpx* (Fig. 1i, j) and *rag1* (Fig. 1k, l) mRNA expression in the CHT at 3 dpf and in the thymus at 4 dpf, respectively, when compared to wild-type zebrafish, indicating impaired definitive haematopoiesis.

Taken together, these findings demonstrate a role of Aqp1a.1 in embryonic haematopoiesis and we next aimed to delineate the mechanisms by which Aqp1a.1-mediated water flux, or cellular hydraulics, regulates haematopoiesis.

### Aqp1a.1 promotes the survival of migrating LPM-derived cells

Endothelial and haematopoietic cells arise from the mesodermal germ layer after gastrulation^1^. Artery-fated angioblasts migrate from the lateral plate mesoderm (LPM) to the embryonic midline between 12 and 18 hpf, where they assemble to generate the DA^11^. We therefore speculated that the reduction in *gata2b^+^*, *runx1^+^* and *cmyb^+^* cells in the DA can result from a decrease in LPM-derived cells reaching the midline.

We first examined the expression of Aqp1a.1 during this early phase of development in a newly generated *TgKI(aqp1a.1-mStayGold)* zebrafish line, which reports endogenous Aqp1a.1 protein localisation, crossed to *Tg(fli1:H2B-mCherry)* zebrafish, which enables visualisation of LPM-derived cells. By performing timelapse confocal imaging, we detected Aqp1a.1 expression that increases from 16 hpf at the plasma membrane of *ffi1^+^* cells located at the LPM (Supplementary Fig. 2 and Movie 1), suggesting a role of cellular hydraulics in regulating the behaviour of *ffi1^+^* cells during migration to the midline. Quantitative analyses of *ffi1^+^* nuclei in *Tg(fli1:H2B-EGFP)* (Fig. 2a) and *aqp1a.1^rk28/rk28^;Tg(fli1:H2B-EGFP)* (Fig. 2b) embryos between 13 and 17 hpf show similar migration speed between wild-type and Aqp1a.1-deficient embryos as cells migrate medially from the LPM to the midline (Fig. 2c, d). However, there is a noticeable difference in nuclear area. In wild-type embryos, there is a continuous increase in nuclear area as *ffi1^+^* cells migrate toward the midline (Fig. 2e). In *aqp1a.1^rk28/rk28^* embryos, nuclear area is significantly larger than wild type between 13 and 14 hpf before decreasing sharply between 14 and 15 hpf (Fig. 2e). This decrease coincided with significant fragmentation and disappearance of *ffi1^+^* nuclei, which we refer to as nuclear fragmentation events, in *aqp1a.1^rk28/rk28^* embryos when compared to wild-type embryos (Fig. 2b, f and g; Movie 2). TUNEL staining revealed widespread cell death in mutant embryos but did not consistently overlap with fragmented *ffi1*^+^ nuclei (Supplementary Fig. 3a, b), indicating that these events are not uniformly associated with TUNEL-detectable DNA fragmentation. Because the cell death mechanism remains unresolved, we use nuclear fragmentation as a morphological readout of cell death. The increased frequency of cell death consequently led to a significant reduction in *gata1a^+^* erythroid progenitors at 12 and 24 hpf (Supplementary Fig. 4a - c), *hbbe3^+^* primitive erythrocytes at 24 hpf (Supplementary Fig. 4d, e), *ffi1^+^* ECs between 14 and 17 hpf (Fig. 2h) and within the DA at 30 and 48 hpf (Fig. 2i, j) in *aqp1a.1^rk28/rk28^* embryos.

**Figure 2.**
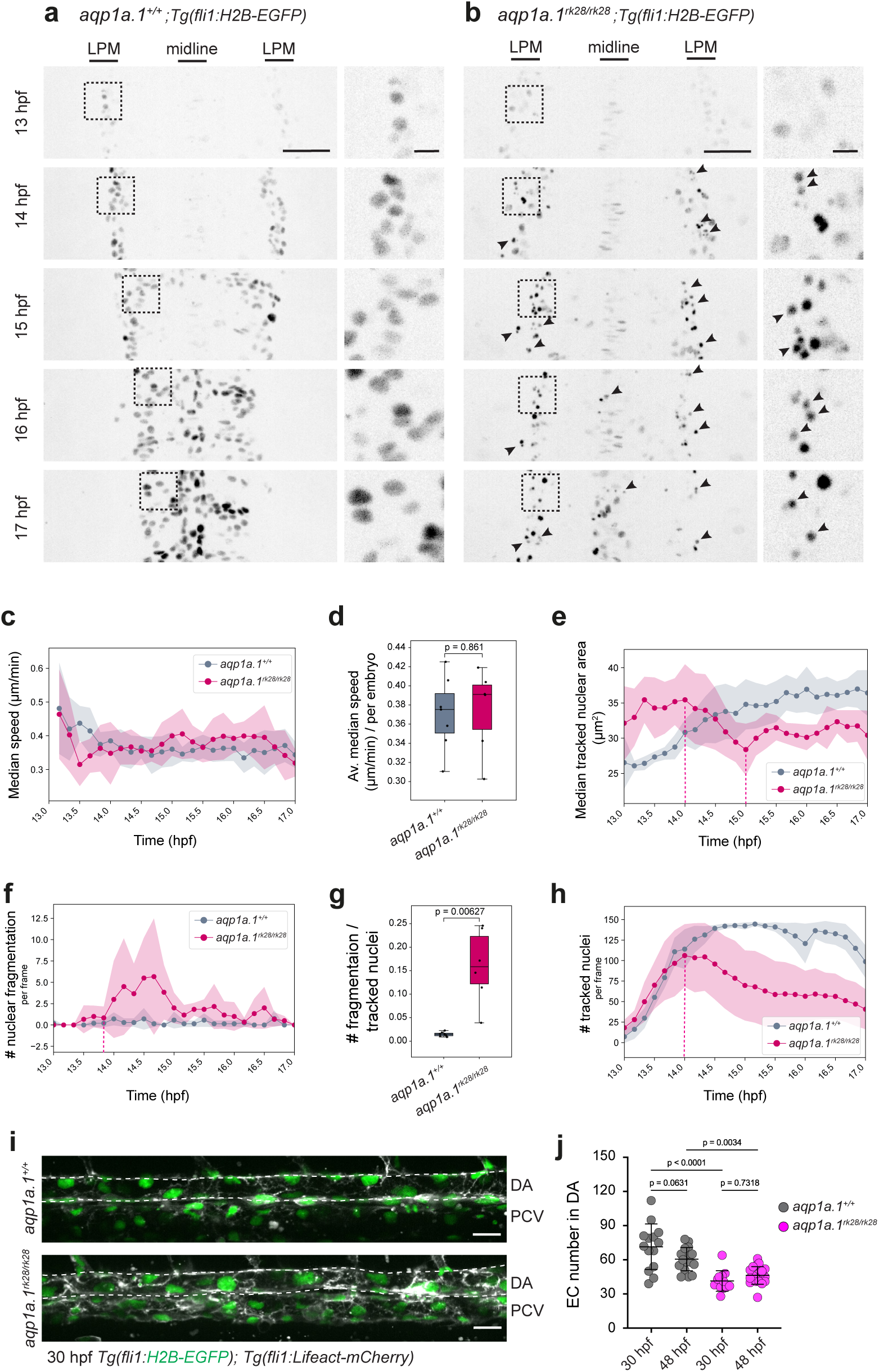
The survival of LPM-derived cells depends on cellular hydraulics. **a**, **b** Still images of LPM-derived *fli1^+^* cells migrating toward the embryonic midline (arrows) in *aqp1a.1^+/+^* (**a**) and *aqp1a.1^rk28/rk28^* (**b**) embryos in *Tg(fli1:H2B-EGFP)* background. A magnified view of the boxed region is shown to the right. Black arrowheads, fragmented nuclei. Time-lapse movies were taken from 13 to 17 hpf. **c**-**h** Analysis of LPM-derived *fli1^+^* cells between 13 to 17 hpf in *aqp1a.1^+/+^* and *aqp1a.1^rk28/rk28^* embryos in *Tg(fli1:H2B-EGFP)* background. Quantification of nuclear velocity (**c**, **d**), area (**e**), nuclear fragmentation (**f**, **g**) and number (**h**) of *fli1^+^*cells in *aqp1a.1^+/+^* (n=7 embryos, 4 independent experiments) and *aqp1a.1^rk28/rk28^* (n=6 embryos, 3 independent experiments) embryos. **d**, **g** two-tailed Welch’s t-test on per-embryo values and exact p values are shown. **i** Representative maximum intensity projection images of *aqp1a.1^+/+^* and *aqp1a.1^rk28/rk28^*embryos in *Tg(fli1:H2B-EGFP);Tg(fli1:Lifeact-mCherry)* transgenic background at 30 hpf. **j** Quantification of EC number in the DA of *aqp1a.1^+/+^* and *aqp1a.1^rk28/rk28^*embryos at 29-32 hpf (*aqp1a.1^+/+^*, n=16; *aqp1a.1^rk28/rk28^*, n=25 embryos) and 48-52 hpf (*aqp1a.1^+/+^*, n=17; *aqp1a.1^rk28/rk28^*, n=24 embryos). Except for **c**-**h**, data were collected from two independent experiments and presented as mean ± SD. Statistical significance was determined by ordinary one-way ANOVA with Sidak’s multiple comparisons test (**j**). Scale bars, 10 μm (**a, c**, dashed boxes), 20 μm (**i**) and 50 μm (**a, b**). DA, dorsal aorta; EC, endothelial cell; LPM, lateral plate mesoderm; PCV, posterior cardinal vein.

These findings demonstrate that Aqp1a.1 promotes the survival of LPM-derived angioblasts during their migration to the embryonic midline, thereby maintaining the endothelial population available for subsequent DA formation and haemogenic specification.

### Aqp1a.1 regulates EC redistribution to the ventral floor of the dorsal aorta

HSPC formation depends not only on establishment of the DA but also on subsequent endothelial morphogenetic events that drive HEC extrusion from the DA. Following the assembly of the DA, ECs redistribute from the dorsal and lateral walls of the DA to the ventral floor, where HECs accumulate before undergoing extrusion ^25,28^. We therefore asked whether Aqp1a.1 contributes to this later morphogenetic process.

We first determined whether Aqp1a.1 is expressed in ECs during this stage by examining its expression and subcellular localisation in the DA. At 24 hpf, approximately 75% of *aqp1a.1*-positive ECs within the DA co-express *runx1* mRNA (Fig. 3a), indicating that Aqp1a.1 is expressed in HECs before extrusion. Aqp1a.1 protein expression was confirmed in ECs in the dorsal and lateral regions of the DA in *TgKI(aqp1a.1-mStayGold)^rk3C^*;*Tg(ffi1:H2B-mCherry)* embryos (Fig. 3b) and in elongated and rounded HECs located at the VDA in *TgKI(aqp1a.1-mScarlet3-H)^rk37^*;*Tg(gata2b:GAL4FF);Tg(UAS:EGFP)* embryos (Fig. 3c) at 50 hpf. Using *TgKI(aqp1a.1-mStayGold)^rk3C^*;*Tg(kdrl:Hsa.HRAS-mCherry)* embryos, we found that Aqp1a.1 is predominantly localised to the plasma membrane (Supplementary Fig. 5a) and remains membrane-associated throughout extrusion including during cell division, invagination and rounding (Fig. 3d and Movie 3). Following extrusion, Aqp1a.1 persists at the plasma membrane of nascent HSPCs where Aqp1a.1 is frequently enriched at their leading edge of dynamic membrane protrusions (Supplementary Fig. 5b and Movie 4).

**Figure 3.**
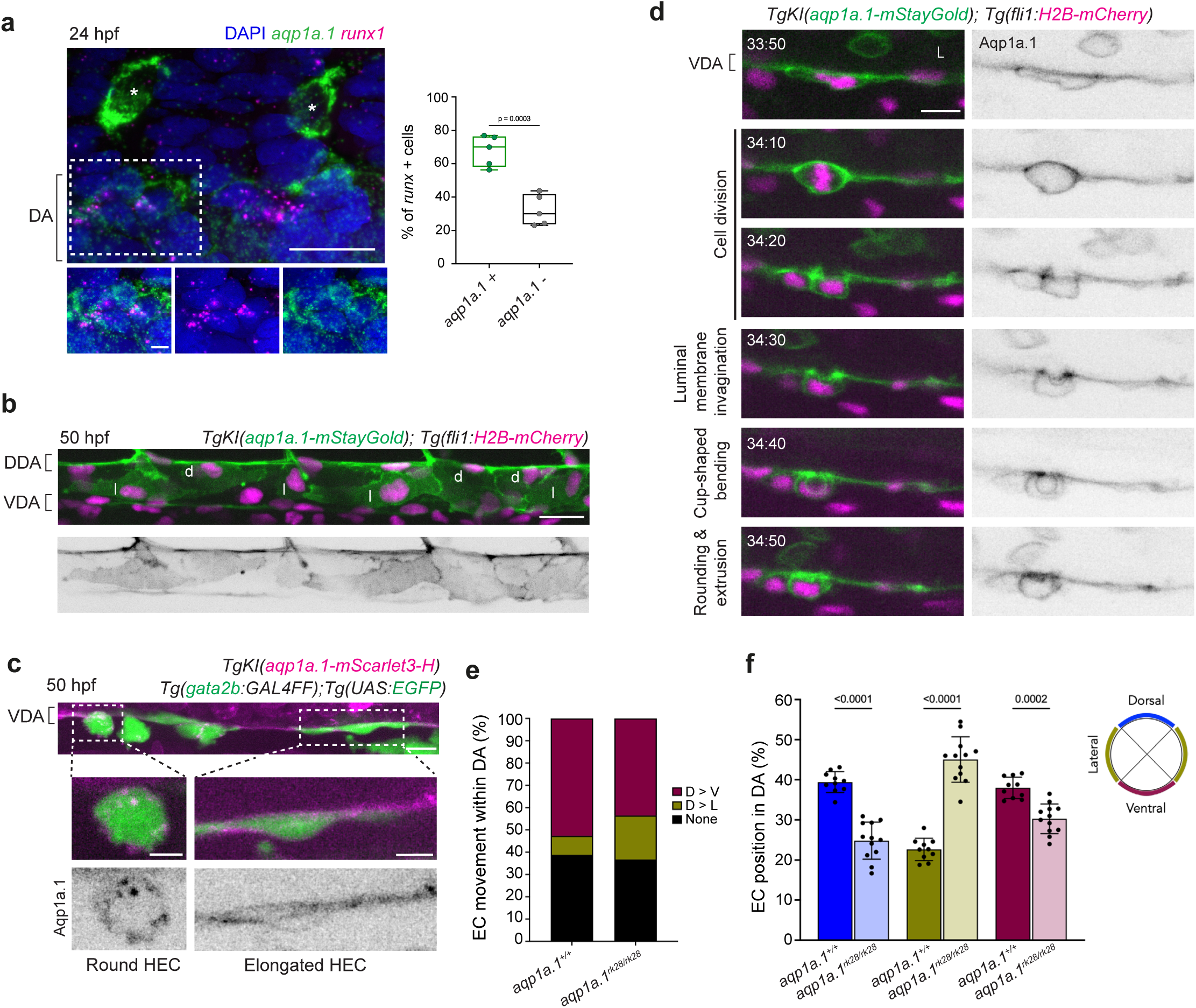
Aqp1a.1 regulates EC redistribution to the ventral floor of the dorsal aorta. **a** Detection of *aqp1a.1* and *runx1* mRNAs by RNAscope *in situ* hybridization in the DA at 24 hpf. Representative maximum intensity projection of confocal z-stacks. A magnified view of the boxed region is shown below. Graph shows percentage of ECs in the DA co-expressing *aqp1a.1* and *runx1* mRNAs (n=5 embryos). *, endothelial tip cells. Scale bar, 10 μm (dashed box) and 20 μm. **b** Heterogeneous expression of Aqp1a.1 protein in ECs of the DA. Representative maximum intensity projection of a *TgKI(aqp1a.1-mStayGold);Tg(fli1:H2B-mCherry)* embryo at 50 hpf. DDA, dorsal wall of the DA; VDA, ventral floor of the DA; d, dorsal EC; l, lateral EC. Scale bar, 20 μm. **c-d** Aqp1a.1 protein is localised at the plasma membrane of HECs during EHT. Maximum intensity projection image of *Tg(gata2b:GAL4FF);Tg(UAS:EGFP);TgKI(aqp1a.1-mScarlet3-H)* embryo at 50 hpf. Magnified views of different HEC shapes are shown below (**c**). Single z-planes extracted from time-lapse movie of *TgKI(aqp1a.1-mStayGold);Tg(fli1:H2B-mCherry)* embryo from 30 to 46 hpf (**d**). L, lumen. Arrow, luminal membrane. Time, hour:minutes. Scale bars, 5 μm (dashed box) and 10 μm. **e** Quantification of EC movement within the DA in *aqp1a.1^+/+^* (n=10) and *aqp1a.1^rk28/rk28^*(n=12) embryos in *Tg(fli1:H2B-EGFP);Tg(fli1:Lifeact-mCherry)* background from 30 to 48 hpf. **f** Quantification of EC position in the DA of *aqp1a.1^+/+^*(n=10) and *aqp1a.1^rk28/rk28^* (n=12) embryos at 48 hpf. Data were collected from 4 independent experiments and presented as mean ± SD. Statistical significance was determined by ordinary one-way ANOVA with Sidak’s multiple comparisons test.

Because Aqp1a.1 is expressed in ECs before HEC extrusion, we next asked whether it contributes to endothelial redistribution within the dorsal aorta. Tracking of ECs located at the dorsal and lateral regions of the DA between 30 and 48 hpf revealed reduced dorsal-to-ventral endothelial movement in *aqp1a.1^rk28/rk28^* embryos when compared to wild-type embryos, leading to abnormal accumulation of ECs within the lateral wall (Fig. 3e, f). Thus, Aqp1a.1 supports both the early survival of artery-fated angioblasts and the subsequent redistribution of ECs toward the ventral floor. Defects in these processes reduce the pool of ECs available for haemogenic specification and EHT, providing a mechanistic explanation for the reduced numbers of HECs and HSPCs observed in *aqp1a.1* mutants (Fig. 1a–d).

### Haemogenic endothelial cell volume reduction during EHT depends on Aqp1a.1-mediated water efflux

The persistent plasma membrane localisation of Aqp1a.1 throughout HEC rounding and invagination further suggests that Aqp1a.1 may function directly during HEC extrusion. Timelapse imaging of *gata2b^+^* cells at the VDA shows the progressive transformation of HECs from an elongated to a rounded morphology (Fig. 4a, upper panel and Movie 5), consistent with previous reports ^25^. During extrusion, HECs form dynamic membrane protrusions and frequently exhibit membrane blebs (Fig. 4a; lower panel and Movie 5), suggesting that these morphological changes are accompanied by elevated intracellular pressure. Because aquaporins regulate cell volume and shape through transmembrane water flux ^30,35,41–43^, we asked whether Aqp1a.1 contributes to cell shape changes that accompany EHT.

**Figure 4.**
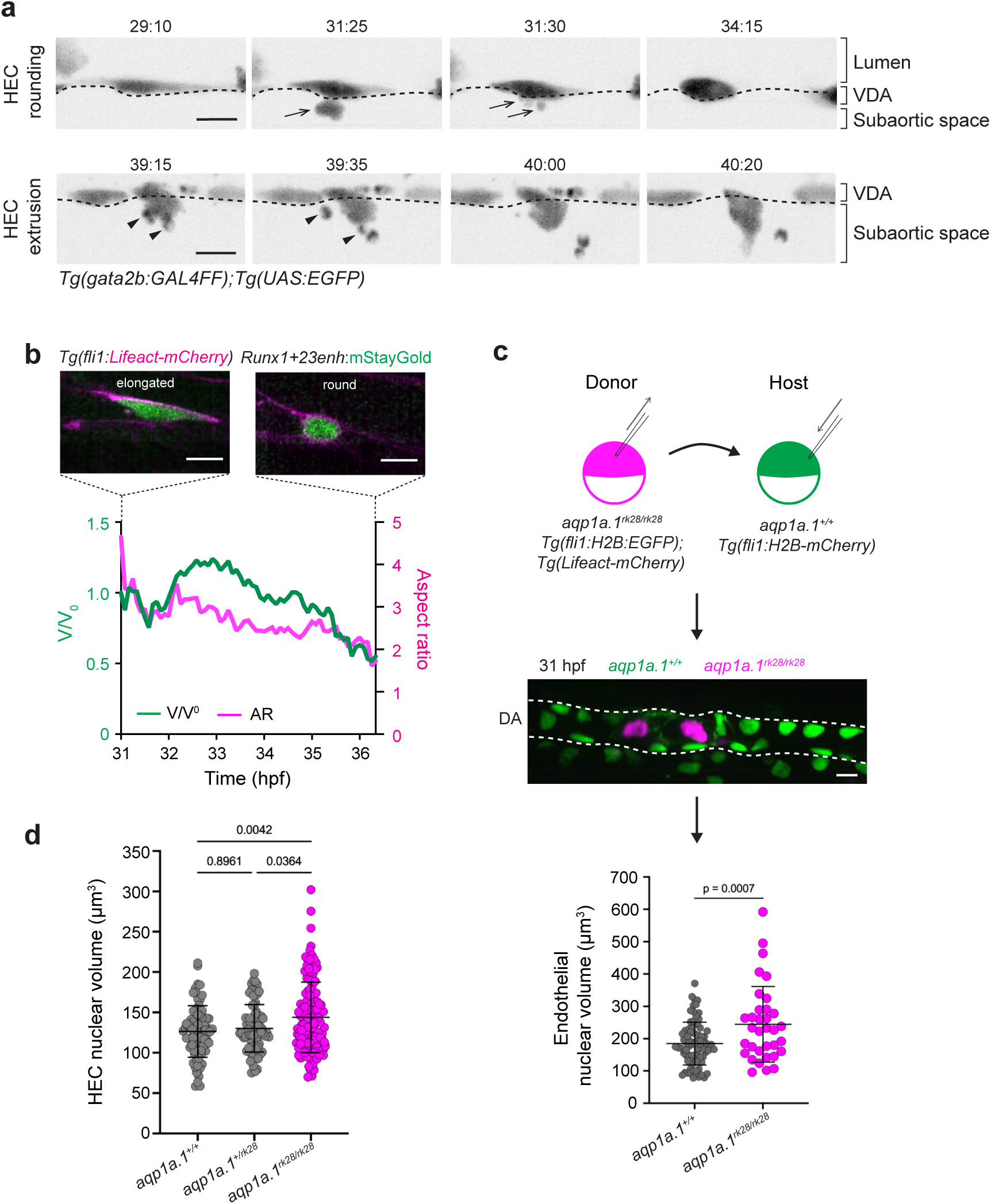
Aqp1a.1 promotes HEC volume decrease during EHT. **a** Cell shape transition of HEC during EHT. Still images from time-lapse movie of HEC dynamics at ventral dorsal aorta (VDA, serrated line) during EHT. Movie of *Tg(gata2b:GAL4FF);Tg(UAS:EGFP)* embryo was taken from 28 to 44 hpf. HEC, hemogenic endothelial cell. Arrows, blebs that retract; arrowheads, blebs that detach. Scale bar, 10 μm. **b** Cell volume decreases as HEC undergoes rounding. Representative graph shows decrease in cell volume and cell aspect ratio of a single cell from 31 to 37 hpf. HEC was labelled by injection of *Runx1+23enh:mStayGold* plasmid into *Tg(fli1:Lifeact-mCherry)* embryo. Deconvolved images (single z-planes) of labelled HEC are shown. Scale bar, 10 μm. **c** Cell transplantation experiments were performed as shown in scheme and representative maximum intensity projection image of 31 hpf embryo is shown. Scale bar, 10 μm. Graph shows quantification of nuclear volume of *aqp1a.1^+/+^* and *aqp1a.1^rk28/rk28^*cells (*aqp1a.1^+/+^*, n=82 nuclei; *aqp1a.1^rk28/rk28^*, n=34 nuclei from 6 embryos). Data are collected from 3 independent experiments and presented as mean ± SD. Statistical significance was determined by two-tailed unpaired *t*-test. **d** Quantification of HEC nuclear volume in *aqp1a.1^+/+^*, *aqp1a.1^+/rk28^*and *aqp1a.1^rk28/rk28^* embryos in *Tg(gata2b:GAL4FF);Tg(UAS:EGFP);Tg(fli1:H2B-mCherry)* background at 30-32 hpf (*aqp1a.1^+/+^*, n=75 nuclei from 11 embryos; *aqp1a.1^+/rk28^*, n=75 nuclei from 10 embryos; *aqp1a.1^rk28/rk28^*, n=4 129 nuclei from 18 embryos). Data are collected from 4 independent experiments and presented as mean ± SD. Statistical significance was determined by ordinary one-way ANOVA with Sidak’s multiple comparisons test.

Tracking and 4D analysis of individual HECs labelled with mStayGold (*Runx1+23enh*:*mStayGold*) at the VDA in wild-type embryos revealed a ∼47% and ∼49% decrease in cell volume and cell aspect ratio, respectively, between 31 to 37 hpf (Fig. 4b and Supplementary Fig. 6a-d), indicating that HECs become smaller as they round up. Tracked cells did not undergo cell proliferation, indicating that the loss of cell volume during HEC extrusion is independent of cell division. This analysis therefore shows that HECs lose volume as they transform from an elongated to round morphology. In contrast, mosaically *kdrl:NLS-TagBFP-P2A-mStayGold* labelled ECs at the dorsal, lateral and ventral regions of the DA as well as extruded cells in *aqp1a.1^rk28/rk28^*;*Tg(fli1:Lifeact-mCherry)* embryos show significant increases in volume when compared to wild type between 30 and 33 hpf (Supplementary Fig. 7a, b).

As nuclear volume and nuclear aspect ratio directly scale with cell volume (Supplementary Fig. 7c, Pearson’s r = 0.8448) and cell aspect ratio (Supplementary Fig. 7d, Pearson’s r = 0.6309), respectively, we also used nuclear volume as a readout of HEC volume to examine HEC volume at a later period of EHT. Between 48 and 52 hpf, wild-type *Tg(fli1:H2B-EGFP);Tg(fli1:Lifeact-mCherry)* embryos demonstrate a spatial difference in EC volume in the DA -nuclear volume decreases by ∼50% from lateral-to-ventral-to-extruded (Supplementary Fig. 7e), supporting the decrease in cell volume during EHT as observed from timelapse imaging (Fig. 4b). While *aqp1a.1^rk28/rk28^* embryos maintain spatial differences in nuclear volume, the loss of Aqp1a.1 significantly increases the nuclear volume of extruded cells and ECs located at the dorsal, lateral and ventral regions of the DA compared to wild-type embryos (Supplementary Fig. 7e). Importantly, in chimaeric wild-type *Tg(fli1:H2B-mCherry)* embryos transplanted with cells from *aqp1a.1^rk28/rk28^*;*Tg(fli1:H2B-EGFP)*;*Tg(fli1:Lifeact-mCherry)* embryos, the nuclear volume of *aqp1a.1^rk28/rk28^* ECs is significantly larger than that of surrounding wild-type ECs (*p* = 0.0007, Fig. 4c), demonstrating a cell-autonomous function of Aqp1a.1 in controlling cell volume in the DA. Finally, to confirm that HEC volume is deregulated, we analysed *gata2b^+^* cells located at the VDA and found that nuclear volume of *aqp1a.1^rk28/rk28^* HECs increased by 13.8% when compared to wild-type embryos (*p* = 0.0364, Fig. 4d).

Together, our findings identify a function of Aqp1a.1 at multiple stages of HSPC development. During early development, Aqp1a.1 promotes the survival of migrating angioblasts to ensure sufficient ECs to form the DA. Within the DA, it regulates the rearrangement of ECs, facilitating their accumulation at the VDA. During EHT, Aqp1a.1 mediates water efflux to reduce HEC volume.

### Piezo channels couple haemodynamic forces to calcium signalling and haemogenic cell swelling

Because ion flux generates osmotic gradients that drive water movement, we next asked which ion-permeable channels and transporters could regulate cell volume during EHT. Piezo1 is a mechanosensitive, non-selective cation channel that mediates epithelial cell extrusion in response to crowding and stretch ^44,45^. Moreover, *piezo1* and *piezo2a* are expressed in the DA between 24 and 58 hpf ^46^ (Supplementary Fig. 8a), prompting us to examine their roles during EHT.

We first examined the spatial distribution of Piezo activity within the DA, using Ca²⁺ activity as its functional readout. In *Tg(fli1:GAL4FF);Tg(UAS:GCaMP7a)* zebrafish ^47^, GCaMP7a activity is consistently higher at the ventral wall (VDA) than in the dorsal region of the DA (DDA) (Supplementary Fig. 9a and Movie 6), yielding VDA/DDA GCaMP7a ratios greater than 1 at both 30 hpf (median 3.0) and 48 hpf (median 4.1), and show no significant difference between the two stages (Supplementary Fig. 9b). Thus, Ca²⁺ activity—a readout of Piezo activity—is spatially enriched in the ventral domain where EHT occurs.

We next resolved Ca²⁺ activity in individual ventral ECs, classified as elongated or round according to their morphology (Fig. 5a–d; Supplementary Fig. 9g–j and Movies 7, 8). Activation of Piezo1 with 20 µM Yoda1 increased mean GCaMP7a intensity, measured as the average F/F₀ during the 20 min following treatment, in both cell classes at 30 and 48 hpf (elongated cells: 1.4-and 1.6-fold, Fig. 5e; round cells: 1.5-and 1.3-fold, Fig. 5f, respectively; *p* ≤ 0.018). Yoda1 also increased Ca²⁺ event frequency by 4.7–8.2 events per cell over 20 min (*p* ≤ 0.004; Fig. 5g, h). By contrast, inhibiting Piezo activity with 1 µM GsMTx4 did not detectably alter either parameter in either cell class or at either stage (*p* ≥ 0.27; Fig. 5e–h and Supplementary Fig. 9c–f, k–n). These findings show that Piezo1 activation is sufficient to enhance Ca²⁺ activity in both elongated and round ventral ECs.

**Figure 5.**
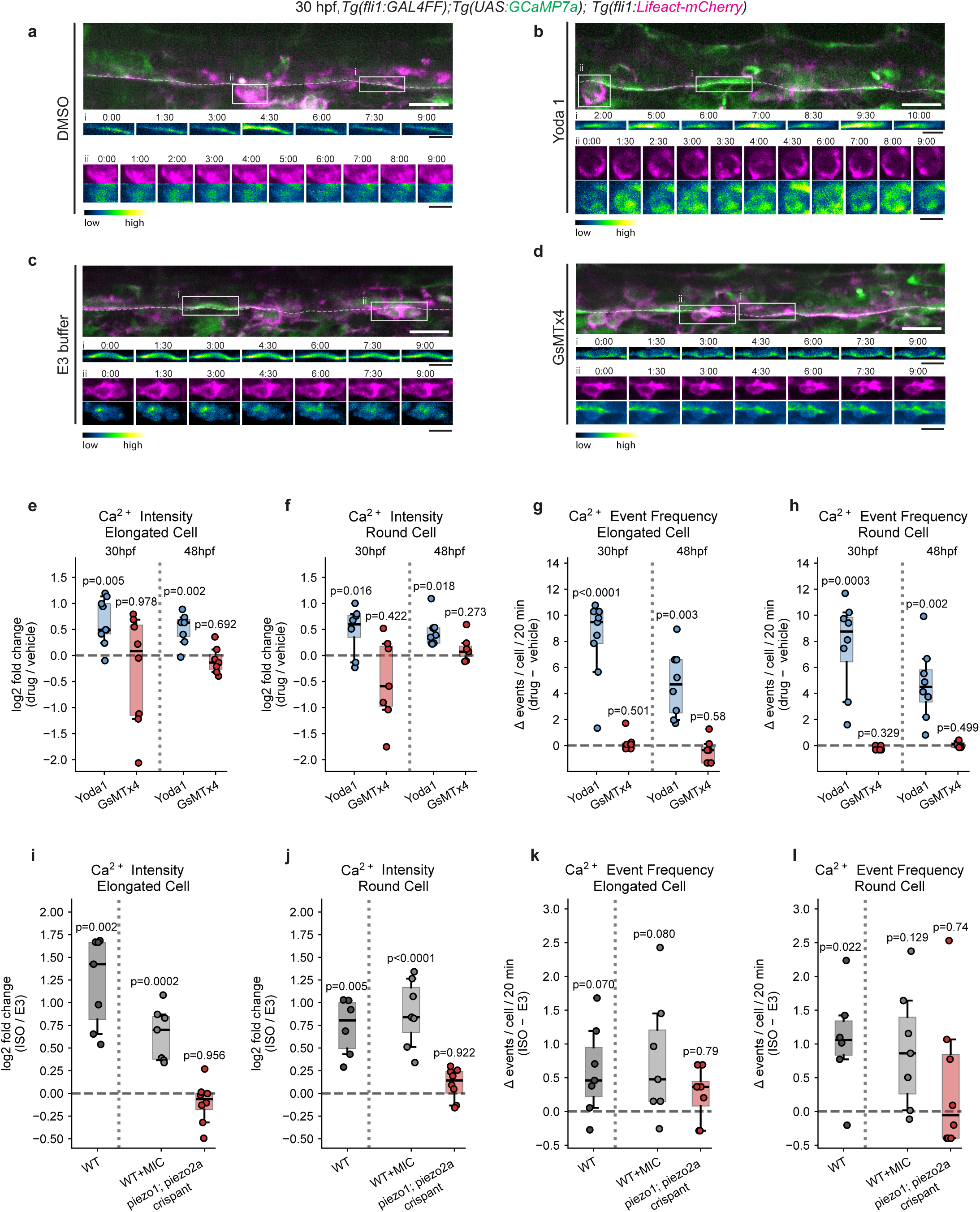
Piezo channels couple haemodynamic forces to calcium signalling. **a**-**d** Representative maximum intensity projections of VDA in *Tg(fli1:GAL4FF); Tg(UAS:GCaMP7a);Tg(fli1:Lifeact-mCherry)* embryos at 30 hpf after treatment with 0.7% DMSO (**a**), 20 µM Yoda1 (**b**), E3 buffer (**c**) or 1 µM GsMTx4 (**d**). GCaMP7a, green; Lifeact-mCherry, magenta; dashed line marks the ventral wall. Boxed regions show elongated (i) and round (ii) ventral EC, classified by morphology and enlarged at bottom. Pseudocolour heat map (LUT: Green Fire Blue) represents GCaMP7a intensity. Scale bars, 20 µm; insets, 10 µm. **e**–**h** Ca²⁺ responses to Piezo1 modulation at 30 and 48 hpf. log2 fold change in mean GCaMP7a intensity (**e**, elongated; **f**, round) and change in Ca²⁺ event frequency (**g**, elongated; **h**, round). Mean GCaMP7a intensity is F/F₀ averaged over the 20 min post-treatment window, where F₀ is the median GCaMP7a signal for that cell during the 5 min pre-treatment recording. Event frequency is expressed as Δ events/cell/20 min. Each dot is one embryo; dashed line indicates no change relative to the vehicle group mean. Data are from 4 independent experiments: 30 hpf (DMSO, n= 23/14 elongated/round cells from 7/6 embryos; Yoda1, n=29/20 from 9/8; E3, n=20/15 from 7/7; GsMTx4, n=21/16 from 8/7) and 48 hpf (DMSO, n=20/23 from 7; Yoda1, n=28/38 from 8; E3, n=15/24 from 6; GsMTx4, n=16/29 from 7 embryos). Statistical significance was determined by two-sided Welch’s t-test. **i**–**l** Ca²⁺ responses in 48 hpf wild-type embryos, mock-injected controls (MIC) and *piezo1;piezo2a* crispants treated with 100 µM isoprenaline hydrochloride (ISO) for 1 hour. log2 fold change in mean GCaMP7a intensity (**i**, elongated; **j**, round) and Δ Ca²⁺ event frequency (**k**, elongated; **l**, round). Data are from four independent experiments in wild-type (E3, n=13/17 elongated/round cells from 6 embryos; ISO, n=18/22 from 7/6) and two in the injected groups (MIC E3, n=23/29 from 7 embryos; MIC ISO, n=20/31 from 7; crispant E3, n=21/36 from 8; crispant ISO, n=21/28 from 8). Statistical significance was determined by two-sided Welch’s t-test (wild-type embryos) and Type II two-way ANOVA (genotype × treatment) followed by Tukey’s HSD (injected groups). Per-group values for every condition are given in Supplementary Fig. 9 and 10.

Because Piezo1 can be activated by blood flow ^48,49^, we next used Ca²⁺ activity as a readout to determine whether haemodynamic forces activate Piezo channels in ventral ECs (Supplementary Fig. 10a–c and Movie 9). Elevating cardiac output with 100 µM isoprenaline hydrochloride (ISO), a β-adrenergic agonist, raised mean GCaMP7a intensity by 2.7-fold in elongated cells and 1.8-fold in round cells (Fig. 5i-j and Supplementary Fig. 10d, e). ISO also increased Ca²⁺ event frequency in round cells, with a non-significant increase in elongated cells (Fig. 5k, l and Supplementary Fig. 10f, g). However, no significant differences were found in both measurements after arresting the heartbeat with 50 mM 2,3-butanedione monoxime (BDM, Supplementary Fig. 10d–g). Importantly, CRISPR/Cas9-mediated depletion of *piezo1* and *piezo2a* (Supplementary Fig. 10h–k and Movie 10) abolished the ISO-induced increase in mean GCaMP7a intensity in both cell classes (Fig. 5i, j and Supplementary Fig. 10l, m), whereas event frequency remained unchanged (Fig. 5k, l and Supplementary Fig. 10n, o). These findings establish Ca²⁺ activity as a functional readout of Piezo activity in ventral ECs and demonstrate that increased haemodynamic forces activate Piezo-dependent Ca²⁺ signalling.

As Piezo1 and elevations in intracellular cations can regulate cell volume in multiple cell types ^50,51^, we next examined whether Piezo activity controls HEC volume during EHT. *Tg(gata2b:GAL4FF);Tg(UAS:EGFP);Tg(fli1:H2B-mCherry)* embryos were treated with Yoda1 or subjected to CRISPR/Cas9-mediated depletion of *piezo1* and *piezo2a*, and the nuclear volume of *gata2b^+^* HECs in the VDA was measured at 30 – 32 hpf. Yoda1 treatment from 25 hpf increased HEC volume by 16% (*p* < 0.0001) whereas depletion of *piezo1* and *piezo2a* reduced nuclear volume by 5.4% (*p* = 0.0223) relative to their respective controls (Fig. 6a, b). Together, these findings indicate that haemodynamic forces activate Piezo-dependent Ca²⁺ signalling in ventral ECs and that Piezo activity promotes HEC swelling.

**Figure 6.**
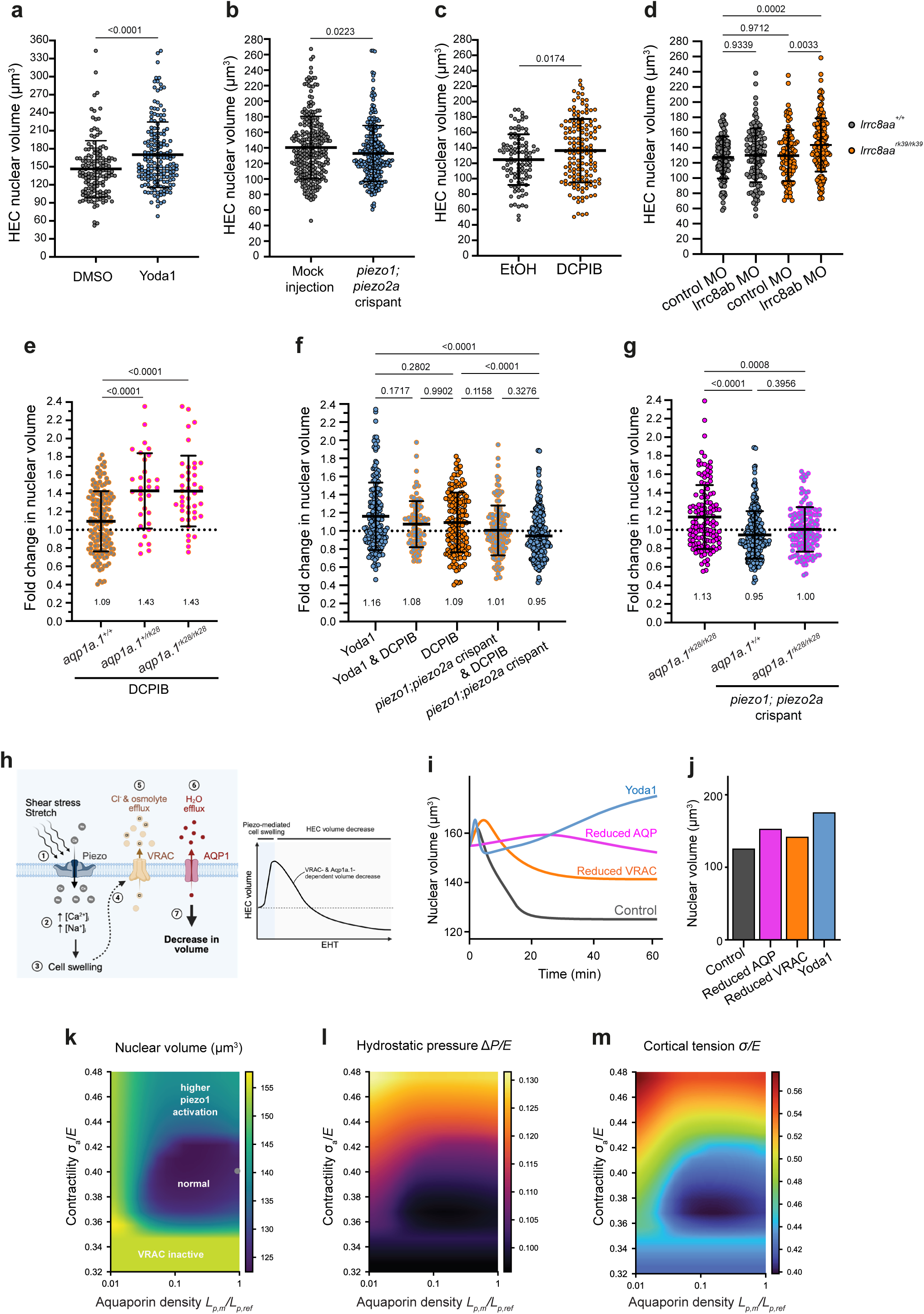
Piezo functions upstream of VRAC-and Aqp1a.1-mediated HEC volume decrease. *Tg(gata2b:GAL4FF);Tg(UAS:EGFP);Tg(fli1:H2B-mCherry)* embryos of the indicated genotypes were treated from 25 to 30 hpf and HEC nuclei volume was quantified at 30-32 hpf. **a** Piezo1 activation in wild-type embryos treated with 0.7% DMSO or 20 µM Yoda1 (n=139 nuclei/13 embryos and n=170/13, respectively). **b** Genetic depletion of *piezo1* and *piezo2a* (mock-injected, n=255 nuclei/20 embryos; crispant, n=268/20). **c** Pharmacological VRAC inhibition (0.05% ethanol, n=102 nuclei/9 embryos and 5 µM DCPIB, n=146/18). **d** Genetic depletion of VRAC (*lrrc8aa^+/+^;*control MO, n=119 nuclei/16 embryos; *lrrc8aa^+/+^;lrrc8ab* MO, n=115/15; *lrrc8aa^rk39/rk39^*;control MO, n=104/10; *lrrc8aa^rk39/rk39^*;*lrrc8ab* MO, n=159/18). **e** Fold change in HEC nuclear volume following DCPIB treatment of *aqp1a.1^+/+^* (n=146 nuclei), *aqp1a.1^+/rk28^* (n=31) and *aqp1a.1^rk28/rk28^* (n=41). **f** Fold change in HEC nuclear volume following individual or combined Piezo and VRAC perturbation in wild-type embryos (Yoda1, n=170 nuclei; Yoda1 plus DCPIB, n=96; DCPIB, n=146; *piezo1;piezo2a* crispant plus DCPIB, n=128; crispant, n=268). **g** Fold change following Piezo depletion in *aqp1a.1^+/+^* and *aqp1a.1^rk28/rk28^* embryos (*aqp1a.1^rk28/rk28^*, n=128 nuclei; *aqp1a.1^+/+^;piezo1;piezo2a* crispant, n=268; *aqp1a.1^rk28/rk28^*;*piezo1;piezo2a* crispant, n=136). Data were collected from two (**a**, **c**-**f**) or three (**b**, **g**) independent experiments and presented as mean ± SD. Statistical significance was determined by two-tailed unpaired *t*-test (**a**-**c**) or ordinary one-way ANOVA with Sidak’s multiple comparisons test (**d**-**g**). **h** Proposed mechanism: haemodynamic forces activate Piezo in VDA endothelial cells (1), increasing intracellular cations (2) and HEC volume (3). Swelling activates VRAC (4), promoting Cl^-^and osmolyte efflux (5), coupled to water efflux through Aqp1a.1 (6), thereby reducing HEC volume (7). **i-m** Mathematical modelling of how contractility, aquaporins, Piezo1 and VRAC regulate cell volume, hydrostatic pressure and cortical tension. Reducing aquaporins (*L_p_*_,*m*_/ *L_p_*_,*ref*_ = 0.01), VRAC (*P_vr_*/*P_vr_*_,*ref*_ = 0.04) or sustaining piezo1 opening with Yoda1 (*α_pz_* = 0.185) cause an increase in cell volume (**i**, **j**). Heatmap of predicted cell volume as a function of cortical contractility and aquaporin density (**k**). Corresponding heatmaps of cytoplasmic hydrostatic pressure (**l**) and cortical tension (**m**). All data obtained 60 mins after change in cell state from reference values.

### VRAC and Aqp1a.1 coordinate osmolyte and water efflux to reduce HEC volume

Although Piezo activity augments HEC volume, HECs undergo a progressive reduction in volume during EHT (Fig. 4a). We therefore sought to identify the mechanism responsible for this volume decrease. ECs of the DA express *lrrc8aa* and *lrrc8ab*^46^ (Supplementary Fig. 8a), which encode the essential Lrrc8 subunits of volume-regulated anion channel (VRAC). Notably, *lrrc8aa* is also detected in the haemogenic endothelium (Supplementary Fig. 8b). VRAC is activated by cell swelling and facilitates the efflux of Cl⁻ and other organic osmolytes, generating an osmotic gradient that drives water efflux during regulatory volume decrease (RVD) ^52–54^. We therefore examined whether VRAC cooperates with Aqp1a.1 to reduce HEC volume during EHT.

Pharmacological inhibition of VRAC activity with 5 µM DCPIB from 25 hpf increased HEC nuclear volume by 9.4% at 30 - 32 hpf when compared to controls (*p* = 0.0174, Fig. 6c). Similarly, genetic depletion of *lrrc8aa* and *lrrc8ab* by injecting an *lrrc8ab* morpholino in *lrrc8aa^rk^*^39/rk39^ embryos significantly increased HEC volume when compared to control morpholino-injected siblings (*p* = 0.0033, Fig. 6d), confirming that VRAC promotes HEC volume reduction during EHT. We further examined the functional interaction between VRAC and Aqp1a.1 by inhibiting VRAC activity with DCPIB in *aqp1a.1* mutant embryos. Whereas VRAC inhibition increased HEC nuclear volume by approximately 1.1-fold in wild-type embryos, its inhibition in *aqp1a.1^+/rk28^* and *aqp1a.1^rk28/rk28^* embryos resulted in a significantly greater increase, reaching approximately 1.4-fold relative to vehicle-treated wild-type embryos (Fig. 6e). Thus, impairment of water permeability sensitises HECs to disruption of osmolyte efflux, indicating that VRAC and Aqp1a.1 cooperate to regulate HEC volume. Interestingly, loss of both *aqp1a.1* alleles did not further increase HEC volume beyond that observed in heterozygous embryos, suggesting that HEC swelling reaches a threshold imposed by the mechanical properties of the cell.

To test more directly whether VRAC and AQP1 constitute an endothelial RVD mechanism, we exposed human aortic endothelial cells (HAECs) to hypotonic shock. Cell volume initially increased, peaking within 2 - 4 min, and subsequently decreased, demonstrating an RVD response (Supplementary Fig. 11c). AQP1 knockdown using siRNA (Supplementary Fig. 11a, b) or VRAC inhibition with DCPIB impaired RVD, leaving cells enlarged at 30 minutes compared to controls (Supplementary Fig. 11d, e). DCPIB also enhanced initial swelling, whereas combined AQP1 knockdown attenuated this effect and impaired RVD to a similar extent as AQP1 knockdown alone (Supplementary Fig. 11f). These findings support coordinated roles for VRAC-mediated osmolyte efflux and AQP1-mediated water efflux in endothelial RVD.

Together, the zebrafish and HAEC experiments support a conserved endothelial volume-regulatory mechanism in which VRAC-mediated efflux of Cl^-^and organic osmolytes lowers intracellular osmotic pressure, while Aqp1a.1/AQP1 provides the membrane water permeability required for the ensuing water efflux. During EHT, this coordinated volume reduction machinery enables HECs to reduce their volume.

### Piezo channels function upstream of HEC volume decrease

Having established that VRAC and Aqp1a.1 compose the downstream machinery responsible for HEC volume reduction, we next asked what initiates this pathway during EHT. Because VRAC is activated by cell swelling^53–55^ and Piezo activation increases HEC volume (Fig. 6a), we hypothesised that Piezo-induced cell swelling acts upstream of VRAC activation to initiate volume reduction.

To test this model, we modulated Piezo activity while simultaneously inhibiting VRAC with DCPIB. Piezo activation with Yoda1 is expected to promote HEC swelling, whereas VRAC inhibition should prevent the ensuing volume decrease. Consistent with this hypothesis, HEC nuclear volume in embryos co-treated with Yoda1 and DCPIB remained comparable to that in embryos treated with Yoda1 alone (Fig. 6f), indicating that Piezo-induced volume expansion is maintained in the absence of VRAC activity. Conversely, the depletion of *piezo1* and *piezo2a* expression, which reduces HEC volume by 5%, should limit volume-dependent activation of VRAC. Accordingly, DCPIB treatment failed to increase HEC volume in *piezo1;piezo2a* crispants (Fig. 6f). While DCPIB treatment caused a 1.09-fold increase in wild-type embryos, combined Piezo depletion and VRAC inhibition resulted in only a 1.01-fold change. Thus, depletion of Piezo suppresses the volume increase caused by VRAC inhibition. Together, these epistasis experiments support a model in which Piezo-mediated cell swelling acts upstream of VRAC-dependent HEC volume decrease during EHT.

Having established that Aqp1a.1 and VRAC are complementary components of HEC volume decrease, Aqp1a.1 should also function downstream of Piezo-mediated cell swelling. If this model is correct, reducing Piezo activity should suppress the enlarged HEC phenotype of *aqp1a.1* mutants (Fig. 4d) by preventing the initial increase in HEC volume that normally triggers volume decrease. To test this prediction, we depleted *piezo1* and *piezo2a* in *aqp1a.1^rk28/rk28^* embryos. Loss of Piezo activity abolished the increase in HEC nuclear volume observed in *aqp1a.1^rk28/rk28^* embryos, restoring nuclear volume to control levels (Fig. 6g). These results therefore place Aqp1a.1 downstream of Piezo-mediated cell swelling and support a model in which Piezo-dependent increases in HEC volume initiate the osmotic changes that drive Aqp1a.1-dependent water efflux during EHT.

Altogether, our findings establish a functional hierarchy in which Piezo channels promote HEC swelling, which subsequently activates VRAC and Aqp1a.1-mediated water efflux to decrease cell volume (Fig. 6h).

### Water retention elevates intracellular hydrostatic pressure and cortical tension

To further understand this coupling of ion flux, water movement and cell volume in HECs, we developed a mathematical model to describe the mechanisms of cell size control. Building on our previous work on multicellular volume regulation ^56^ and growth ^57^, we considered how the change in cell volume over time is described by a competition between osmotic and hydrostatic pressure gradients across the membrane. Cellular osmotic pressure evolves primarily from electrochemically-driven ion transport across the membrane through active pumps, passive leak channels, and mechanosensitive pathways (e.g. Piezo1, VRAC), and also from cytoplasmic biomolecules (see Methods for details). Our model supports a pathway whereby stimulated Piezo1 promotes a cation influx to swell the cell above a threshold for VRAC activation. A subsequent loss of anions and other osmolytes through VRAC drives a reduction in cell fluid volume through aquaporins (Supplementary Fig. 12). To simulate the elevated actomyosin contractility during HEC rounding and extrusion ^24^, we increased contractility from baseline conditions. Upon associated activation of piezo1 and VRAC downstream, our analysis suggests that cells with a high aquaporin density will rapidly reduce in volume (Fig. 6i-k, control). Such contractility also increases both hydrostatic pressure (Fig. 6l) and cortical tension (Fig. 6m), compounding the loss of water. Interestingly, suppression of aquaporins, VRAC activity, or Piezo1 activation with Yoda1 are all predicted to increase cell volume (Fig 6i-k; Supplementary Fig. 12) by magnitudes in direct agreement with our experimental data. Reducing VRAC expression directly lowers the rate of anion and osmolyte loss, and maintains a higher cell volume. With Yoda1 treatment, there is sustained Piezo1 opening in a mechanism distinct from physical stimulation, facilitating a continuous cation influx which can offset reductions in osmotic pressure from VRAC. In cases where there is aquaporin depletion, the rate of water loss across the membrane is significantly impaired and cell volume becomes less sensitive to contractility and osmotic pressure (Fig. 6h-j). Our model suggests that these cells cannot shrink as rapidly to relieve hydrostatic pressure (Fig. 6l) or cortical tension (Fig. 6m). With such elevated and sustained cortical tension, cells are at higher risk of membrane rupture arising from mechanisms such as cytoskeletal detachment and blebbing.

### Osmolyte and water efflux promotes HEC robustness during cell shape transition

To test these predictions, we performed timelapse imaging of *aqp1a.1^rk28/rk28^*;*Tg(fli1:H2B-EGFP)*;*Tg(fli1:Lifeact-mCherry)* embryos to examine the behaviours of Aqp1a.1-deficient cells during their extrusion from the VDA. Similar to our observation during the migration of *ffi1^+^* cells from the LPM to the midline, ectopic cell death occurred between 30 and 48 hpf predominantly at the VDA, leaving behind fragmented nuclei (Supplementary Fig. 13a-c and Movie 11). This phenotype resembled the abortive EHT previously observed in Runx1 morphants ^10^. Time-lapse imaging of *aqp1a.1^rk28/rk28^*; *Tg(gata2b:GAL4FF);Tg(UAS:EGFP)* embryos over the same period confirmed that the dying ventral ECs are HECs, as indicated by the fragmentation of *gata2b^+^* cells (Fig. 7a, b and Movie 12). Analysis of chimaeric embryos, in which *aqp1a.1^rk28/rk28^* cells were transplanted into wild-type embryos, revealed a higher frequency of cell death in Aqp1a.1-deficient ECs in the DA than among wild-type ECs (Fig. 7c, d), demonstrating a cell-autonomous requirement for Aqp1a.1 in EC survival. We next tested the model’s prediction that reducing contractility would protect Aqp1a.1-deficient HECs from the mechanical consequences of impaired water efflux. Treatment with the ROCK inhibitor Y-27632, which reduces actomyosin contractility, rescued the ectopic cell-death phenotype in *aqp1a.1^rk28/rk28^* embryos (Fig. 7b). This result supports a mechanism in which Aqp1a.1-mediated water efflux relieves contractility-induced mechanical stress, thereby preserving HEC integrity and survival during cell shape transitions.

**Figure 7.**
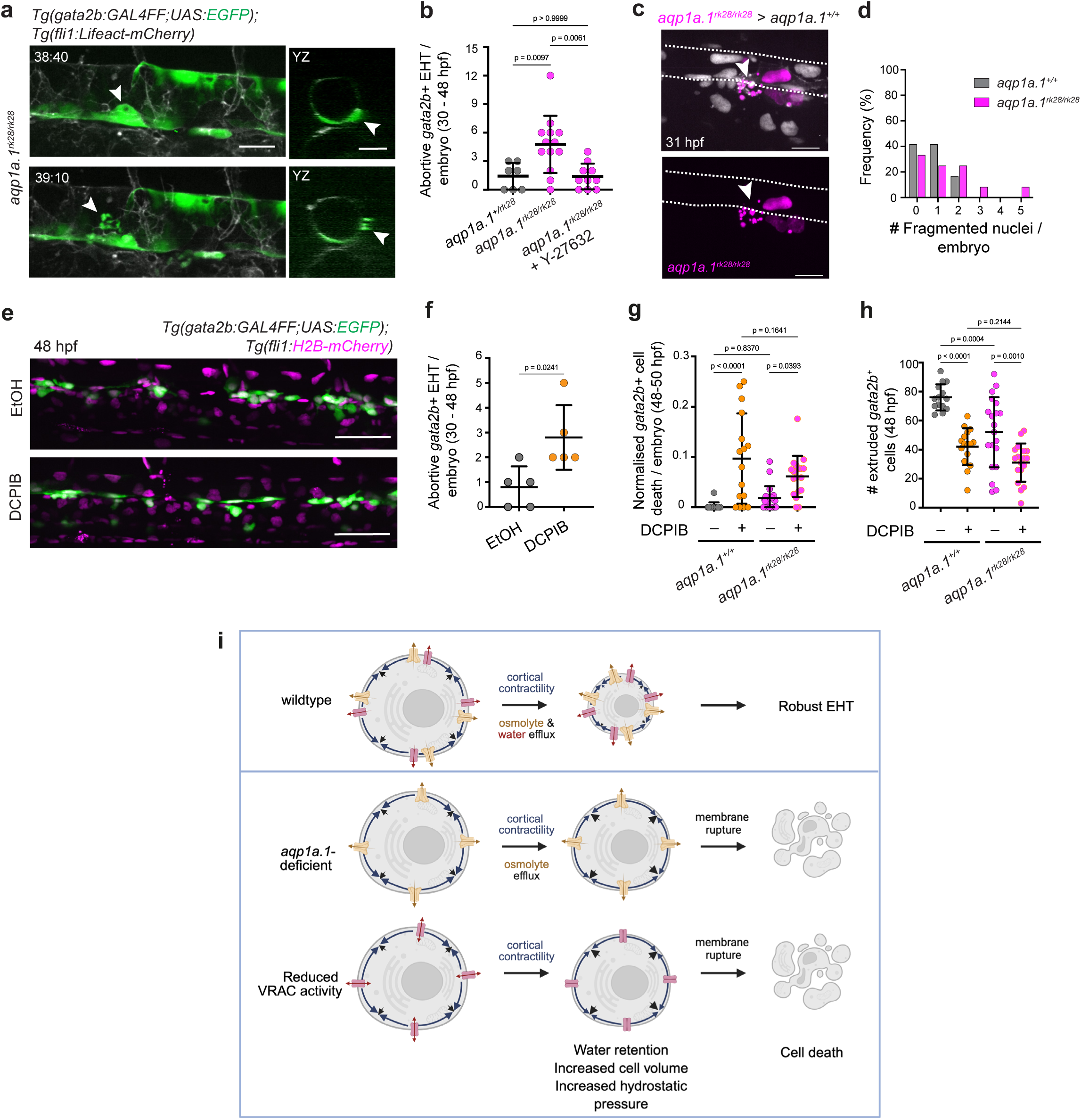
Disrupted osmo-hydraulics results in enhanced cell death. **a–d** Aqp1a.1-depleted HECs undergo increased abortive EHT. Still images from a time-lapse movie of *aqp1a.1^rk28/rk28^*;*Tg(gata2b:GAL4FF);Tg(UAS:EGFP);Tg(fli1:Lifeact-mCherry)* embryo (**a**). Arrowhead, fragmented HEC at VDA. Quantification of *gata2b^+^* cells that underwent abortive EHT between 30 to 48 hpf in *aqp1a.1^+/rk28^*, *aqp1a.1^rk28/rk28^* and *aqp1a.1^rk28/rk28^* embryos treated with 100 μM Y-27632 (**b**, *aqp1a.1^+/rk28^*, n=7 embryos; *aqp1a.1^rk28/rk28^*, n=11 embryos; Y-27632 treated *aqp1a.1^rk28/rk28^*, n=10 embryos from 3 independent experiments). Representative maximum intensity projection image of chimaeric *aqp1a.1^+/+^*embryo with transplanted *aqp1a.1^rk28/rk28^* cells at 31 hpf showing abortive EHT (**c**, arrowhead). Serrated lines mark DA boundaries. The frequency of nuclear fragmentation in host *aqp1a.1^+/+^* and transplanted *aqp1a.1^rk28/rk28^*cells (**d**, n=12 embryos). Data were collected from 3 independent experiments. **e-h** Inhibition of VRAC increases HEC death during EHT and decreases HSPC extrusion. Representative maximum intensity projection of *Tg(gata2b:GAL4FF); Tg(UAS:EGFP); Tg(fli1:H2B-mCherry)* embryos at 48 hpf (**e**). *aqp1a.1^+/+^*and *aqp1a.1^rk28/rk28^* embryos in *Tg(gata2b:GAL4FF);Tg(UAS:EGFP);Tg(fli1:H2B-mCherry)* background were treated with 0.05% ethanol or 5 μM DCPIB between 30 and 48 hpf (**g**, **h**). Number of *gata2b^+^*cells that underwent abortive EHT events was quantified between 30 and 48 hpf (**f**, ethanol, n=5 *aqp1a.1^+/+^* embryos; DCPIB, n=5 *aqp1a.1^+/+^* embryos from two independent experiments) and at 48 hpf (**g**, ethanol, n=14 *aqp1a.1^+/+^*and 20 *aqp1a.1^rk28/rk28^* embryos; DCPIB, n=16 *aqp1a.1^+/+^*and 18 *aqp1a.1^rk28/rk28^* embryos from two independent experiments). Results are presented as mean ± SD. Statistical significance was determined by Brown-Forsythe and Welch ANOVA with Dunnett’s T3 multiple comparisons test (**b**), two-tailed unpaired *t*-test with Welch’s correction (**f**) and ordinary one-way ANOVA with Sidak’s multiple comparisons test (**g**, **h**). **i** VRAC and Aqp1a.1 coordinate osmolyte and water efflux to reduce cell volume, relieve intracellular pressure and preserve HEC integrity during EHT. Scale bar, 15 μm (**c**), 20 μm (**a**) and 50 μm (**e**).

As the inhibition of Cl^-^and osmolyte flux through VRAC also increased EC size at the VDA, we next examined whether VRAC inhibition similarly compromises HEC survival. Time-lapse imaging of DCPIB-treated *Tg(gata2b:GAL4FF);Tg(UAS:EGFP)* embryos from 30 to 48 hpf showed increased fragmentation of *gata2b*^+^ cells relative to control embryos (Fig. 7e – g). This was accompanied by a significant decrease in the number of extruded *gata2b*^+^ cells at the VDA at 48 - 52 hpf in DCPIB-treated embryos (Fig. 7h) and in *lrrc8aa^rk^*^39/rk39^ embryos injected with *lrrc8ab* morpholino (Supplementary Fig. 14), demonstrating that VRAC activity also promotes HEC survival and successful extrusion during EHT. Importantly, combined inhibition of VRAC and Aqp1a.1 activity further compromised HEC survival. DCPIB treatment of *aqp1a.1^rk28/rk28^* embryos significantly increased the number of fragmented *gata2b^+^* cells when compared to untreated *aqp1a.1^rk28/rk28^* embryos (Fig. 7g). This elevated cell death was accompanied by a further reduction in the number of *gata2b*^+^ cells at the VDA (Fig. 7h). These findings indicate that VRAC-mediated osmolyte efflux and Aqp1a.1-mediated water efflux cooperate to preserve HEC integrity during the pronounced cell shape and volume changes associated with EHT.

In sum, we conclude that disruption of ion and osmolyte transport and cellular hydraulics compromises HEC survival during cell rounding and extrusion in EHT (Fig. 7i), thereby diminishing HSPC formation and definitive haematopoiesis.

## Discussion

Most studies of EHT have focused on transcriptional and mechanotransductive programs that govern HEC specification and HSPC formation, whereas the biophysical mechanisms underlying HEC morphological transformation remain less well understood. Here, we identify an osmo-hydraulic volume-regulation mechanism that enables HECs to withstand contractile forces associated with HEC rounding and extrusion. Our findings support a pathway in which haemodynamic forces activate Piezo to promote HEC swelling, which subsequently triggers VRAC-mediated osmolyte efflux and Aqp1a.1-mediated water efflux. This coordinated response reduces HEC volume, preserves cell integrity and supports definitive haematopoiesis.

Pulsatile blood flow imposes shear stress and circumferential stretch on ECs, with ECs lining the VDA experiencing greater circumferential stretch than those in the dorsal and lateral regions of the DA ^25,58^. Pulsation-mediated stretch was recently shown to activate Piezo1 to promote long-term haematopoietic stem cell formation in mouse embryos and human pluripotent stem cell-derived models^58^, but the downstream cellular mechanisms remained unclear. Our live imaging shows that increased haemodynamic forces elevate Ca²⁺ activity in ventral ECs and that this response depends on Piezo1 and Piezo2a. Pharmacological activation of Piezo1 also increases Ca²⁺ activity and HEC volume, whereas depletion of *piezo1* and *piezo2a* reduces HEC volume. This swelling likely results from the osmotic consequences of Piezo-mediated cation influx, including Na⁺ and Ca²⁺ entry. Genetic and pharmacological epistasis further places Piezo-dependent swelling upstream of VRAC and Aqp1a.1. Piezo depletion suppresses the volume increase caused by VRAC inhibition and restores the enlarged HECs of *aqp1a.1* mutants to control volume. These findings extend the established role of Piezo in HSPC formation by linking haemodynamic mechanosensing to HEC volume regulation.

We propose that Piezo-dependent swelling initiates a regulatory volume decrease (RVD) response. RVD typically involves the efflux of K⁺, Cl⁻ and organic osmolytes, followed by osmotically driven water loss ^50,52^. In HECs, VRAC mediates Cl⁻ and osmolyte efflux, generating an osmotic gradient that drives water efflux through Aqp1a.1. Accordingly, pharmacological or genetic disruption of VRAC increases HEC volume, and this increase is exacerbated when Aqp1a.1-mediated water permeability is impaired. Inhibition of VRAC or loss of Aqp1a.1 also increases cell death during EHT, while mathematical modelling predicts that osmolyte and water efflux limit the hydrostatic pressure and cortical tension generated during HEC contraction. Thus, as actomyosin contractility drives HEC rounding and extrusion, Aqp1a.1 may function as a pressure-relief valve that prevents intracellular forces from exceeding the mechanical tolerance of the cell. Aqp1a.1 loss similarly causes swelling and nuclear fragmentation in LPM-derived cells migrating towards the embryonic midline, suggesting that volume regulation protects cells during multiple force-generating morphogenetic processes. However, aquaporin loss does not universally induce cell death: endothelial tip cells lacking Aqp1a.1 and Aqp8a.1 survive during zebrafish sprouting angiogenesis ^30^, while aquaporins promote apoptosis in some other cell types ^59,60^. The consequences of altered water permeability therefore depend on cellular state and mechanical context.

Emerging evidence indicates that cell volume and water flux not only maintain cellular homeostasis but can also influence cell fate and differentiation. For example, volume regulation has been implicated in the differentiation of human epidermal keratinocytes and mesenchymal stem cells^61,62^. These findings raise the possibility that impaired water transport contributes to the reduction in haematopoiesis in Aqp1a.1-deficient embryos by altering differentiation of LPM progenitors into endothelial or blood lineages. However, expression of haemangioblast marker *tal1* and angioblast marker *etsrp* is preserved at 12 hpf (Supplementary Fig. 15a, b), arguing against a broad defect in initial progenitor specification. Together with the increased loss of LPM-derived cells and HECs, our results suggest that the reductions in erythroid progenitors, primitive erythrocytes, ECs, HECs and HSPCs arise predominantly from compromised cell survival.

Our study also reveals distinct, stage-dependent functions of Aqp1a.1 in HSPC development. Aqp1a.1 first promotes the survival of LPM-derived progenitors, thereby maintaining sufficient ECs for DA formation. It subsequently facilitates endothelial redistribution towards the VDA and, during EHT, mediates water efflux to reduce HEC volume. Our findings also differ mechanistically from those of Sato et al., who demonstrated that aquaporins promote vacuole formation and HEC rounding during avian EHT^27^. We do not consistently observe intracellular vacuoles in rounding or extruding zebrafish HECs. The occasional vacuolar structures arise through apical membrane invagination and fusion during HEC extrusion from the VDA and subsequently disappear (Fig. 3d). Moreover, loss of Aqp1a.1 increases HEC volume without altering the aspect ratio of ECs at the VDA (Supplementary Fig. 16). Aquaporins may therefore support EHT through different cellular mechanisms depending on species and whether HSPCs emerge as intra-aortic clusters or as individual cells extruding into the subaortic space. Nevertheless, the requirement for aquaporins in both avian and zebrafish EHT suggests that cellular hydraulics is conserved in this transition.

A limitation of our study is that we do not directly measure intracellular osmolarity, ion flux, hydrostatic pressure or cortical tension. Quantitative ion imaging, electrophysiology and direction measurements of cellular mechanics will be required to define how ion and water transport are coupled during EHT. Although we establish a role for Piezo and VRAC in HEC volume regulation, they have additional functions that may also contribute to HEC behaviour. Piezo-mediated cation influx can alter membrane potential and downstream pathways regulating cytoskeletal organisation and gene expression ^63,64^, whereas VRAC transports organic osmolytes including taurine, glutamate and myo-inositol in addition to Cl^-^and can regulate membrane potential and extracellular signalling ^52,65^. Other ion channels and transporters involved in regulatory volume decrease such as K^+^/Cl^-^cotransporters and calcium-activated chloride channels may also contribute to HEC behaviour ^52^. Thus, although our epistasis experiments establish a functional hierarchy between Piezo, VRAC and Aqp1a.1, additional transport and signalling pathways may influence endothelial morphogenesis and haemogenic specification.

Together, our findings identify osmo-hydraulics volume regulation as a determinant of HEC survival during EHT. By coupling Piezo1-dependent cell swelling to VRAC-mediated osmolyte efflux and Aqp1a.1-mediated water efflux, HECs balance the forces required for rounding and extrusion against the need to preserve membrane integrity. More broadly, our results suggest that cells undergoing force-generating morphological transitions must actively regulate their volume and intracellular pressure to remain viable during fate transitions. Similar osmo-hydraulic mechanisms may operate in other developmental contexts involving extensive cell shape changes while maintaining cellular integrity and could potentially be harnessed to improve the robustness of in vitro HSPC generation.

## Methods

### Zebrafish maintenance and strains

Zebrafish (*Danio rerio*) were raised and staged according to established protocols^66^. The following published zebrafish lines were used in this study: *aqp1a.1^rk28^* ^30^, *lrrc8aa^rk3S^* (this study), *Tg(fli1:zgc:114046-EGFP)^ncvCS^* ^67^ [referred to as *Tg(fli1:H2B-EGFP)^ncvCS^*], *Tg(fli1:Lifeact-mCherry)^ncv7^* ^68^, *Tg(fli1:myr-mCherry)^ncv1^* ^69^, *Tg(fli1:h2bc1-mCherry)^ncv^*^31^ ^47^ [referred to as *Tg(fli1:H2B-mCherry)^ncv31^*]*, Gt(gSAIz-GAL4FFM)^nkgSAIzGFFM1770A^* ^70^ [a gene trap line containing *GAL4FF* in exon 3 of *gata2b* gene, and hereafter denoted as *gata2b:GAL4FF* line], *Tg(5xUAS:EGFP)^nkuasgfp1a^* ^71^ *, Tg(UAS:GCaMP7a)* ^72^ *, Tg(fli1:GAL4FF)^ubs3^* ^73^ *, TgBAC(pdgfrb:EGFP)^ncv22^* ^74^ *, Tg(kdrl:Hsa.HRAS-mCherry)^sS1C^* ^75^,and *TgKI(aqp1a.1-mStayGold)^rk33^* (this study) and *TgKI(aqp1a.1-mScarlet3-H)^rk37^* (this study). Zebrafish were maintained on a 14h/10h light/dark cycle. Fertilised eggs were collected and raised in E3 medium at 28^0^C. To inhibit pigmentation in embryos older than 24 hpf, 0.003% *N*-Phenylthiourea (PTU) (Sigma Aldrich, P7629) in E3 medium was used. All animal experiments were approved by the Institutional Animal Care and Use Committee at RIKEN Kobe Branch (IACUC).

### Plasmid generation, gene knockdown and microinjections

Plasmids *pDs-kdrl:NLS-TagBFP-P2A-mStayGold* and *pDs-Runx1+23enh:mStayGold* were generated using In-Fusion HD Cloning kit (Takara Bio Inc., Cat# 639650). Plasmids encoding mStayGold ^76^, mScarlet3-H ^77^, the mouse *Runx1 +23* enhancer and the zebrafish *kdrl* promoter were generous gifts from Yasushi Okada (RIKEN BDR, Japan), Kiryl Piatkevich (Westlake University Hangzhou, China), Anne Schmidt (Institut Pasteur, France) and Massimo M. Santoro (University of Turin, Italy), respectively. To avoid unwanted *Tol2* re-transposition from the donor site in transgenic zebrafish lines generated with *Tol2* transposon, we used *Ac/Ds* transposon system from the maize ^78^ for mosaic expression of DNA constructs. Plasmid DNA (10 ng/μl) and *Ac* transposase mRNA (100 ng/μl) were co-injected into one-cell stage zebrafish embryos. *Ac* transposase mRNA was synthesized using the mMESSAGE mMACHINE SP6 kit (Invitrogen, Cat# AM1340) according to manufacturer protocol and purified with RNA Clean and Concentrator-5 kit (Zymo Research, Cat# R1015).

The antisense morpholino oligo (MO) 5’-TATGAACTTGACTTACTCTGTTGC-3’ ^79^, targeting the splice donor site of *lrrc8ab*, and standard control morpholino were purchased from Gene Tools. To knockdown *lrrc8ab*, 1.5 ng of *lrrc8ab* MO was injected at one-cell stage *lrrc8aa^rk3S/rk3S^* and *lrrc8aa^+/+^* embryos in *Tg(gata2b:GAL4FF);Tg(UAS:EGFP);Tg(fli1:H2b-mCherry)* background.

CRISPR interference was used to knockdown *piezo1* and *piezo2a*. Three Alt-R crRNAs per gene were designed and ordered from IDT (Integrated DNA Technologies, USA). For each gene, the three crRNAs were pooled and annealed to Alt-R tracrRNA in IDT duplex buffer by heating to 95^0^C for 5 min and cooling to room temperature. Ribonucleoproteins were assembled by combining the two guide cocktails with Alt-R HiFi Cas9 nuclease V3 (Integrated DNA Technologies, Cat# 1081061), 0.3 M KCl and phenol red, and incubating at 37^0^C for 10 min, followed by microinjection into the cell of one-cell stage embryos. Control embryos from the same clutch were injected in parallel with an identical mix containing tracrRNA and Cas9 but no targeting crRNA. Targeting efficiency was assessed after each experiment by lysing injected and control embryos and amplifying a short fragment spanning each target site; efficient cutting was observed as a smear or multiple bands on a 2% sodium borate agarose gel, compared with a single tight band in controls (see Supplementary Fig. 17a). Larger amplicons spanning each locus were Sanger sequenced to confirm the lesions. Sequences of the primers and crRNAs are listed in Supplementary file 1. To evaluate cutting efficiency at the *piezo1* and *piezo2a* genetic loci *in vivo*, RNPs were injected into *TgBAC(pdgfrb:EGFP);Tg(fli1:myr-mCherry)* embryos, and vascular mural cells were imaged at 5 dpf alongside with uninjected siblings (Supplementary Fig. 17b). EGFP-positive cells associated with the dorsal aorta were counted. Consistent with a requirement for Piezo signalling in mural cell recruitment, *piezo1/piezo2a* crispants showed a reduction in the number of *pdgfrb^+^* mural cells associated with the dorsal aorta at 5 dpf compared with uninjected siblings (Supplemental Fig. 17c), reproducing the loss of *transgelin*-positive vSMCs reported following *piezo1/piezo2a* suppression in zebrafish ^80^.

### Generation of *lrrc8aa* zebrafish mutant line

The *lrrc8aa* mutant line was generated by CRISPR/Cas9 mutagenesis. Briefly, an Alt-R crRNA 5’-CATGACGATGGAAATGTAGT-3’ targeting the second exon of *lrrc8aa* (see Supplementary Fig. 18) was annealed to Alt-R tracrRNA, assembled with Alt-R HiFi Cas9 nuclease V3, and injected into the cell of one-cell stage embryos within 15 min of fertilisation. F_0_ founder fish were identified by outcrossing CRISPR/Cas9-injected fish with wild type fish and screening the offspring for mutations at 2 dpf using Sanger sequencing.

### DNA isolation and genotyping

Genomic DNA was isolated from embryos or fin clips using HotSHOT method ^82^. To screen for mutations, target genomic DNA regions were amplified via PCR. A 550-bp product for the *aqp1a.1* gene was generated using the primers 5’-CGCCTCCAGATTCATTAGCAGGA-3’ and 5’-GTAAGTGAACTGCTGCCAGTGA-3’ and subsequently analyzed by Sanger sequencing with the primer 5’-ATGAACGAGCTGAAGAGCAAG-3’. Similarly, a 2441-bp product of the *lrrc8aa* gene was amplified using the primers 5’-GGAGAAGAAAACACATTGACACGT-3’ and 5’-CATTAGAAAAACCCAACCTGCCC-3’, followed by Sanger sequencing using the primer 5’-CTCATTTTAGCCAGAATCGTGCAG-3.

### cDNA synthesis

Total RNA was isolated from whole 12 hpf (for cloning *etsrp* and *tal1*) and 28 hpf (for cloning *gata1a*, *hbbe3*, *mpx* and *runx1*) zebrafish embryos with TRI Reagent using Direct-zol RNA MicroPrep kit (Zymo Research, Cat# R2061-A) according to manufacturer’s protocol. Total RNA was store at -80^0^C. The first-strand cDNA was synthesized from 1 µg of a total RNA by oligo(dT) priming using the SuperScript III First-Strand synthesis system (Invitrogen, Cat# 18080-051) according to the manufacturer’s protocol. cDNA samples were stored at -20^0^C.

### Generation of TgKI(aqp1a.1-mStayGold) and TgKI(aqp1a.1-mScarlet3-H) zebrafish lines

An ALT-R crRNA targeting the last exon of *aqp1a.1*, 5’-GCCACTGATTATGAGGTCAA-3’, was designed and ordered from IDT (Integrated DNA Technologies, USA). The donor template plasmid was created by inserting 335-bp of genomic sequence, flanking the *aqp1a.1* stop codon, into *pUC1S* using In-Fusion cloning. mStayGold or mScarlet3-H with a short linker Gly-Gly-Ala-Ala-Gly-Gly-Ser-Arg were inserted into the resulting construct, using In-Fusion cloning to produce *pUC1S_aqp1a.1-mStayGold* and *pUC1S_aqp1a.1-mScarlet3-H*. In-Fusion cloning was used to introduce 4 silent single nucleotide changes in the protospacer to produce the final construct *pUC1S_aqp1a.1-mStayGold^ΔPS^* and *pUC1S_aqp1a.1-mScarlet3-H^ΔPS^*. The knock-in donors were produced by amplifying an 842-bp PCR product, using *pUC1S_aqp1a.1-mStayGold^ΔPS^* or *pUC1S_aqp1a.1-mScarlet3-H^ΔPS^* as the template. 0.4 μl of 100 μM ALT-R crRNA, 0.4 μl of 100 μM ALT-R tracrRNA and 0.5 μl of IDT duplex buffer was incubated for 5 min at 95^0^C and then cooled to room temperature to anneal the heteroduplex gRNA. 1 μl of the annealed gRNA was incubated with 1.5 μl 1 M KCl (filter sterilized), 0.5 μl phenol red, 0.3 μl IDT ALT-R HiFi Cas9 nuclease V3, 1.2 μl water and 0.5 μl of donor (250 ng/μl) at 37^0^C for 30 minutes. 1-2 nl of the mix was injected into the one-cell stage embryos within the first 15 mins of fertilisation. Only embryos with broad mStayGold or mScarlet3-H expression were raised. Founders were outcrossed to AB zebrafish, and mStayGold or mScarlet3-H positive embryos were sequenced to check for errors at the repair junctions. Primers used to generate both lines provided in Supplementary file 1.

### Cell transplantation

Cell transplantation was performed as previously described ^81^. Donor cells were harvested from embryos derived from *aqp1a.1^rk28/rk28^;Tg(fli1:H2B-EGFP);Tg(fli1:Lifeact-mCherry)* zebrafish at blastula stages and transplanted into the lateral marginal zone of wild-type *Tg(ffi1:H2B-mCherry)* recipient embryos between 4.5 and 6 hpf. Transplanted embryos were raised under standard conditions and screened for successful incorporation of donor endothelial cells. Chimaeric embryos were imaged between 30 and 32 hpf and nuclei volume from donor and recipient embryos were measured.

### Whole mount RNAscope in situ hybridization

RNAscope^®^ *in situ* hybridization was conducted as previously described ^30^ using RNAscope Multiplex Fluorescent Reagent kit v2 (Advanced Cell Diagnostics, Cat# 323100) and RNAscope probes, *Dr-aqp1a.1* (Advanced Cell Diagnostics, Cat# 893521) and *Dr-runx1* (Advanced Cell Diagnostics, Cat# 1260971-C2). Nuclei were counterstained with DAPI ready-to-use solution overnight at 4^0^C. Prior to imaging, embryos were rinsed in PBST and kept in 70% glycerol in PBS at 4^0^C in the dark. Embryos were deyolked and flat mounted for imaging on an inverted Olympus FV3000 laser scanning confocal microscope (Evident, Japan) using an Olympus UPlanXApo 60x/NA 1.42 oil immersion objective. Image acquisition was controlled by the Olympus software.

### Whole mount chromogenic *in situ* hybridization

Chromogenic *in situ* hybridization was conducted as previously described ^30^. AP-conjugated anti-digoxigenin antibody (Roche, Cat# 11093274910, 1:5000) and NBT (Roche, Cat# 11383213001) and BCIP (Roche, Cat# 11383221001) were used as antibody and substrate for color reaction. Plasmids encoding zebrafish cDNAs of *cmyb* and *rag1* were gift from Shigetomo Fukuhara (Nippon Medical School, Japan). DNA templates for *in situ* probes were generated by PCR amplification from cDNA using a high-fidelity KOD-Plus-Neo DNA polymerase (Toyobo, Cat# KOD-401) (see Supplementary file 1 for primers) and cloned using NEB PCR cloning kit (New England BioLabs, Cat# E1202S). RNA probes for *cmyb, runx1, rag1, mpx, gata1a, hbbe3, etsrp* and *tal1* were *in vitro* synthesized either with MEGAscript SP6 (Invitrogen, Cat# AM1330) or MEGAscript T7 (Invitrogen, Cat# AM1333) kits and digoxigenin-labeled rNTPs (Roche, Cat# 11277073910) and purified using RNA Clean and Concentrator-5 kit (Zymo Research). Image z-stacks were captured by Leica M205FA microscope with digital camera DFC7000T controlled by LAS X software (Leica, Germany), using the same exposure, magnification and illumination setting for each embryo. For quantitative analysis, images were first inverted and converted to 8-bit grayscale. A region of interest (ROI) was manually defined around the expression signal to measure mean pixel intensity. An identical ROI in a non-stained area was used as a background. Pixel intensity was normalised by subtracting the background intensity from the signal intensity.

### Live confocal imaging

Dechorionated embryos were anesthetized using 0.16 mg/mL tricaine (Sigma Aldrich, Cat# E10521) and mounted in 0.8% low-melt agarose (Bio-Rad, Cat# 1613111) in E3 medium containing tricaine and 0.003% phenylthiourea on a glass bottom 35-mm dish (MatTek Corporation, USA). Embryos were imaged using an inverted Olympus IX83-ZDC2 microscope (Olympus, Japan) equipped with a motorised stage and a spinning disc confocal unit (CSU-W1; Yokogawa, Japan) with single laser input (405/488/561/640 nm). Depending on an experimental setup, images were captured with different objectives using Zyla 4.2 sCMOS camera (Andor, Oxford Instruments) with array size 2048×2048 pixels (fixed 1x magnification) and 50 μm pinhole. Image acquisition was controlled by Andor iQ3 software and z-stack images were captured for 3D reconstruction. TagBFP was excited by a 405 nm laser, EGFP and mStayGold were excited by a 488 nm laser, and mCherry and mScarlet3-H were excited by 561 nm laser. Images were processed using Fiji software (NIH, version 2.16.0/1.54p) with brightness and contrast adjustments. Z-stacks of images are presented as maximum intensity projections.

### Pharmacological treatments

DCPIB (Tocris, Cat. No. 1540) was prepared as 10 mM in EtOH and stored at -20^0^C. Yoda1 (Tocris, Cat. No. 5586) was prepared as 2.8 mM solution in DMSO and stored at 4^0^C. GsMTx4 (Fujifilm, Cat. No. 4393-S) was prepared as 5 μM solution in MilliQ water and stored at -20^0^C. ISO (Sigma, Cat. No. I5627-5G) was prepared as 50mM in MilliQ water and stored at -20^0^C. BDM (Sigma, Cat. No. B0753-25G) was prepared as 500mM in MiliQ water and stored at -20^0^C. Y-27632 (Selleck Chemicals, Cat. No. S6390) was prepared as 50 mM in DMSO and stored at -80^0^C. All chemicals were diluted to the desired concentration in E3 medium. For cell and nuclear volume measurement, embryos were treated with 5 μM DCPIB, 20 μM Yoda1 or 1 μM GsMTx4 from 25 to 30 hpf and imaged between 30 and 32 hpf. For quantifying the number of extruded *gata2b^+^* cells, embryos were treated from 30 to 48 hpf and imaged between 48 and 52 hpf.

### TUNEL assay and antibody staining

24 hpf *Tg(fli1:H2B-EGFP)* and *aqp1a.1^rk28/rk28^;Tg(fli1:H2b-EGFP)* embryos were fixed in freshly prepared 4% methanol-free PFA (ThermoFisher Scientific, Cat# 28908) in PBS at 4^0^C overnight and dehydrated with methanol at -20^0^C. Embryos were then rehydrated with PBST (PBS plus 0.01% Tween-20) and permeabilized with 10 μg/ml proteinase K (ThermoFisher Scientific, Cat# EO0491) for 15 min. Embryo were processed with Click-iT Plus TUNEL assay kit (Invitrogen, Cat# C10619) according to manufacturer’s protocol. To stain EC nuclei, embryos were blocked with blocking buffer (1x Blocking reagent, Roche) for 1 hour. The anti-GFP primary antibody (Santa Cruz Biotechnology, Cat# sc-9996) was diluted (1:200) in blocking buffer and Alexa Fluor 488 anti-mouse antibody (Invitrogen, Cat# A-11017, 1:1000 diluted in blocking buffer) was used as secondary antibody. Embryos were deyolked, flat mounted and imaged using an inverted Olympus FV3000 laser scanning confocal microscope (Evident, Japan) with an Olympus UCPlanFLN 20x/NA 0.7 objective.

### Transfection and siRNA-mediated protein knockdown

Human aortic endothelial cells (HAECs) were purchased from Lonza (CC-2535, lot no. 20TL231227) and used between passages 3 and 5. HAECs were cultured in EGM-2 medium (Lonza, CC-3024) supplemented with the EGM-2 BulletKit (Lonza, CC-3124), containing 2% fetal bovine serum (FBS), bovine brain extract, ascorbic acid, human epidermal growth factor, hydrocortisone and antibiotics. Cells were maintained at 37°C in a humidified atmosphere containing 5% CO₂.

For siRNA-mediated knockdown, HAECs were seeded at a density of 7.9 × 10⁴ cells/cm² and transfected with 150 nM siRNA using DharmaFECT 1 transfection reagent (Dharmacon, T-2001-02) in Opti-MEM® Reduced Serum Medium (Gibco, 31985070) without antibiotics. After 24 h, the transfection medium was replaced with EGM-2 supplemented with 2% FBS. Cells were cultured for an additional 48 h before total protein extraction or experimental treatment. ON-TARGETplus Human AQP1 siRNA-SMARTpool (Dharmacon, L-021494-00-0010) was used for AQP1 knockdown, and ON-TARGETplus Non-targeting Pool siRNA (Dharmacon, D-001810-10) was used as the control siRNA. Successive AQP1 knockdown was confirmed with anti-AQP1 antibody immunostaining and Western blotting (see Supplementary Fig. 11a, b).

### Hypotonic shock and drug treatment

For hypotonic shock experiments, cells were exposed to a hypotonic solution prepared by mixing two volumes of Milli-Q water with one volume of EGM-2 medium (2:1, v/v). The hypotonic solution was applied after the baseline imaging period.

To inhibit volume-regulated anion channels (VRACs), cells were pretreated with 5 µM DCPIB (Tocris Bioscience, Cat. No. 1540) for 30 min before hypotonic shock. Ethanol (0.05% final concentration) was used as the vehicle control for DCPIB treatment.

For AQP1-knockdown experiments, control siRNA-and AQP1 siRNA-transfected cells were subjected to hypotonic shock. AQP1-knockdown cells were also pretreated with 5 µM DCPIB for 30 min before hypotonic shock to assess the contribution of VRAC activity to the cellular response to hypotonic stress.

### Live-cell imaging

For live-cell imaging, cells were seeded in 8-well chambered slides (ibidi, Cat. No. 80807) and maintained at 37°C using a temperature-controlled enclosure equipped with a heated stage-top incubator (ibidi, 12720) during imaging. Before imaging, nuclei were stained with NucBlue™ Live ReadyProbes™ Reagent (R37605, Invitrogen) for 3 min and washed with culture medium. For inhibitor experiments, cells were pretreated with the respective inhibitor for 30 min before imaging.

Images were acquired at 3-min intervals for 9 min to establish a baseline. Hypotonic shock was then applied, and imaging was continued at 1-min intervals for 30 min. For AQP1-knockdown experiments, both control siRNA-and AQP1 siRNA-transfected cells were imaged following the same protocol, with or without DCPIB pretreatment as indicated.

Confocal z-stacks were acquired using an inverted Olympus IX83/Yokogawa CSU-W1 spinning-disk confocal microscope equipped with a Zyla 4.2 CMOS camera (Andor) and a Olympus UPlanSApo 40x/NA 1.25 silicon immersion objective (pixel size=0.162 μm) with a z-step interval of 0.16 µm.

### Western blot

Cells were lysed and proteins were denatured in SDS sample loading buffer for 5 min at 95 °C. Proteins were separated on 12% polyacrylamide gels by SDS-PAGE using a Mini-PROTEAN Tetra Cell (Bio-Rad, 1658004JA). The separated proteins were transferred onto 0.45-µm PVDF membranes (Thermo Fisher Scientific, 88518) using a Mini Trans-Blot Electrophoretic Transfer Cell (Bio-Rad, 170-3930) at 100 V for 90 min. Membranes were blocked for 1 h at room temperature in blocking buffer consisting of PBS containing 3% bovine serum albumin (BSA) and 0.05% Tween 20. The membranes were then incubated overnight at 4 °C with primary antibodies against mouse anti-AQP1 (MA1-20214, Thermo Fisher Scientific) and rabbit anti-α-tubulin (ab52866, Abcam), each diluted 1:1,000 in blocking buffer. After washing with PBS containing 0.05% Tween 20 (PBS-T), membranes were incubated with HRP-conjugated goat anti-mouse IgG (H+L) secondary antibody (31430, Thermo Fisher Scientific) for detection of AQP1 and HRP-conjugated goat anti-rabbit IgG (H+L) secondary antibody (31460, Thermo Fisher Scientific) for detection of α-tubulin, each diluted 1:10,000 in blocking buffer, for 1 h at room temperature. Following further washing with PBS-T, chemiluminescent signals were developed using SuperSignal™ West Dura Extended Duration Substrate (37071, Thermo Fisher Scientific) and detected using a chemiluminescence imaging system (LAS-3000mini, Fujifilm).

### Immunofluorescence

HAECs were transfected in 24-well plates and maintained for 48 h. Cells were then reseeded at 0.5 × 10⁴ cells per well in 8-well ibidi chambered slides. After an additional 24 h, cells were fixed with 4% paraformaldehyde (PFA) in PBS (15710, Electron Microscopy Sciences) for 15 min at room temperature, permeabilized with 0.1% Triton X-100 for 1 h and blocked with 1% bovine serum albumin (BSA) for 1 h at room temperature. Cells were incubated with primary antibody against anti-AQP1 (1:100; MA1-20214, Thermo Fisher Scientific) diluted in blocking buffer for 1 h at room temperature. Following three washes, cells were incubated with anti-mouse Alexa Fluor 488 secondary antibody (1:1000; Invitrogen, A11008), for 1 h at room temperature. Nuclei were counterstained with DAPI (14 μM; Thermo Fisher Scientific, 62248).

### Cell and nuclear volume measurements

To measure cell and nuclear volume, single cells in the dorsal aorta of *Tg(fli1:Lifeact-mCherry)* and *aqp1a.1^rk28/rk28^;Tg(fli1:Lifeact-mCherry)* embryos were labelled through mosaic expression of either *pDs-kdrl:NLS-TagBFP-P2A-mStayGold* plasmid (dual labelling of EC nuclear and cytoplasm) or *pDs-Runx1+23enh:mStayGold* plasmid (specific labelling of HECs). Embryos were imaged between 29 and 32 hpf using an Olympus UPlanSApo 60x/NA 1.2 water immersion objective (pixel size=0.108 μm) with an optical z-plane interval of 0.26 μm. For Figure 4e, *Tg(fli1:H2B-EGFP);Tg(fli1:Lifeact-mCherry)* and *aqp1a.1^rk28/rk28^;Tg(fli1:H2B-EGFP);Tg(fli1:Lifeact-mCherry)* embryos were imaged between 48 and 52 hpf using an Olympus UPlanSApo 40x/NA 1.25 silicon immersion objective (pixel size=0.162 μm) with an optical z-plane interval of 1.0 μm. Before analysis, raw images were deconvolved with Huygens Spinning Disk Deconvolution tool (Huygens Essential 22.10 software, Scientific Volume Imaging B.V.), following standard protocol established in the unit (classic MLE deconvolution algorithm, automatic estimation of background, optimized iteration mode with 50 iterations and quality threshold=0.01). Cells and nuclei were subsequently segmented, 3D reconstructed and volume measured using Object Analyzer tool in Huygens Essential software (Scientific Volume Imaging B.V.). The intensity threshold was automatically determined using Otsu method, and seed and sigma were manually adjusted.

### Quantification of EC number in the DA

Measurements were performed in *Tg(fli1:H2B-EGFP);Tg(fli1:Lifeact-mCherry)* and *aqp1a.1^rk28/rk28^;Tg(fli1:H2B-EGFP);Tg(fli1:Lifeact-mCherry)* embryos. Confocal z-stacks images of embryos were taken at 29-32 and 48-52 hpf using an Olympus UPlanSApo 40x/NA 1.25 silicon immersion objective. EC nuclei were counted in the DA region between ISVs no. 7-15 using Fiji software.

### Nuclear tracking and analysis of lateral plate mesoderm cells

To analyse the behaviour of cells arising from the lateral plate mesoderm (LPM), *Tg(fli1:H2B-EGFP)* and *aqp1a.1^rk28/rk28^;Tg(fli1:H2B-EGFP)* embryos were imaged from 13 to 17 hpf with a temporal resolution of 10 minutes using CSU-W1 spinning disk confocal microscopy and an Olympus UCPlanFLN 20x/NA 0.7 objective (pixel size=0.325 μm). Nuclei were segmented with StarDist2D using the pre-trained 2D_versatile_fluo model. Analysis was restricted to the LPM by a region of interest defined automatically from the distribution of nuclear centroids in early frames. Notochord-associated objects were then excluded by locating the embryonic midline as the gap between the two bilateral bands of LPM nuclei: a one-dimensional density profile of nuclear centroid x-positions was computed from nucleus-sized objects in early frames, candidate peak pairs were scored on the depth of the intervening valley and on how completely the pair bracketed the tissue, and the midline was taken as the midpoint of the highest-scoring pair, with its orientation set by a rotation sweep maximising the same criterion. Objects lying within 20 µm of the fitted midline were excluded across the entire time series. Objects of 20-60 µm² were treated as intact LPM nuclei and tracked; objects below 10 µm² were classified as fragments, excluded from tracking but retained for fragmentation detection. Tracking was performed with TrackPy using nearest-neighbour linking in physical units (search radius 8.5 µm, temporal memory 2 frames), after limiting each frame to the 150 largest nuclei and enforcing a minimum inter-nuclear separation of 3 µm. Tracks shorter than three frames were discarded, and remaining track ends were joined across gaps of up to three frames and 10 µm. Nuclear fragmentation was defined as a tracked nucleus of at least 30 µm² whose area fell below 60% of its previous value, accompanied by two to eight fragments appearing within two frames and 8 µm, each no larger than 30% of the pre-fragmentation nuclear area. This last bound excludes mitotic divisions, whose two daughter nuclei are each approximately half the size of the parent. Both within-track fragmentation, in which tracking continued after area collapse, and track ending fragmentation, in which the nucleus disappeared from tracking, were included. Fragmentation frequency was normalised to the number of tracked nuclei. Nuclear area, migration speed, fragmentation fraction, number of tracked nuclei and pre-fragmentation nuclear area were aligned to absolute developmental time (hpf), summarised across embryos and reported as mean ± s.d. Statistical comparisons used two-tailed Welch’s t-tests on per-embryo values, and exact p values are reported without star-based significance annotations. A total of 7 wild type and 6 aqp1a.1 mutant embryos, from 4 and 3 independent experiments respectively, were analysed.

### Calcium imaging and analysis

Confocal imaging was performed on an inverted Olympus IX83 microscope equipped with a Yokogawa CSU-W1 spinning-disk unit, using an Olympus UPLSAPO 30×/NA 1.05 silicone oil objective and a Zyla 4.2 CMOS camera (Andor). Image acquisition was controlled with Andor iQ3. For baseline calcium imaging (“pre”), embryos were imaged for 5 min with a 1 µm z-interval (72 z-stacks) and a 30-second time interval. Embryos were then released from agarose and treated with vehicle (E3 buffer or 0.7% DMSO), 20 µM Yoda1, or 1 µM GsMTx4 for 30 minutes; 100 µM isoprenaline hydrochloride (ISO) for 1 hour; or 50 mM 2,3-butanedione monoxime (BDM) for 5 minutes. After treatment, embryos were remounted in agarose and imaged for an additional 30 minutes (“post”) using identical acquisition settings. During post imaging, embryos were continuously covered with the corresponding drug-containing medium. Each time series was registered in Fiji (ImageJ 1.54p) with Linear Stack Alignment with SIFT (MultiChannel, default settings), and quantitative measurements were exported with a custom Fiji macro (ImageJ 1.54p). ROIs included the ventral region of the dorsal aorta (VDA), dorsal region of the dorsal aorta (DDA), background, and individual endothelial cell ROIs classified as elongated or round based on morphology. For each ROI and timepoint, the macro exported measurements for both channels (GCaMP7a and Lifeact-mCherry), including mean and maximum intensity values, and automatically computed background-subtracted signals (*mean_bgsub*, *max_bgsub*). All downstream analyses were performed using the background-subtracted means. All analysis and plotting were performed directly on these Fiji-exported CSV files in Python (3.10.19).

To assess spatial differences in calcium activity, background-subtracted mean GCaMP signals were used to compute ventral-to-dorsal ratios in the pre-phase:

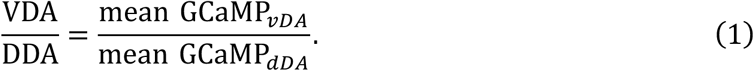

Ratios were calculated per timepoint and summarised for each embryo as the median VDA/DDA ratio, which was visualised using embryo-level boxplots. To correct for potential differences between pre-and post-imaging caused by remounting, a Lifeact-based scale correction was applied at the embryo level using the vDA signal:

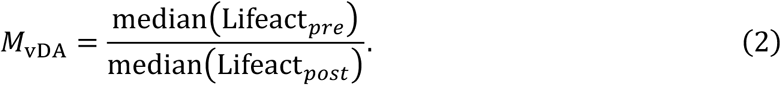

For each embryo, all post-phase GCaMP values were corrected as:

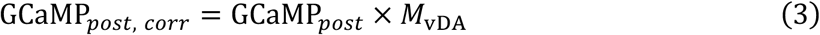

Pre-phase values were not rescaled.

For single-cell analysis, baseline fluorescence *F*_0_was defined for each cell as the median GCaMP signal during the pre-phase:

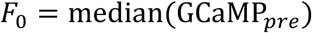

Normalised calcium traces were calculated as:

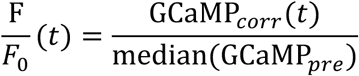

where GCaMP*_pre_* corresponds to uncorrected pre-phase values and GCaMP*_corr_*(*t*) to Lifeact-corrected post-phase values. Single-cell F/F₀ traces were generated for elongated and round ECs. All single-cell metrics were computed on the first 20 min of the post phase. Candidate peaks were required to satisfy at least one of two amplitude-based criteria: (i) a relative increase of at least 10% compared to the larger of the adjacent frames, or (ii) a local prominence criterion within a ±2-frame temporal window, defined relative to the 20th percentile of local F/F values. In addition, peaks were required to exceed a threshold defined on each cell’s own pre-phase as the median plus twice the robust standard deviation (1.4826 × MAD). For each embryo and cell class, event frequency was calculated as the mean number of events per cell per 20 min, and mean GCaMP7a intensity as the mean of F/F_0 over the window, in each case averaging across all analysed cells of that class within the embryo; the embryo is the unit of replication. Fold changes in the figures are a display transform; tests used the per-embryo values. Yoda1 and GsMTx4 were each compared with their own vehicle (0.7% DMSO and E3 buffer) by two-sided Welch’s t-test, uncorrected, as the comparisons use separate vehicle groups. Isoprenaline was compared with E3 buffer alone by two-sided Welch’s t-test; where BDM was included as a third group, the three conditions were compared by one-way ANOVA followed by pairwise Welch’s t-tests with Holm-Bonferroni correction. Mock-injected controls and *piezo1;piezo2a* crispants were compared by Type II two-way ANOVA (genotype × treatment) with Tukey’s HSD. Exact p values are reported in the plots.

### Mathematical modelling of cellular hydromechanics

Changes in cell volume are regulated by the flux of ions that can be transported across the cell membrane. Considering a concentration of internal permeable ions *N*_j_ of species *j*, the associated osmotic pressure is given by the van’t Hoff relation such that Π*_p_* = ∑ *N*_j_ *RT*/*V* where *R* is the gas constant, *T* is the absolute temperature, and *V* is the cell volume. Cell volume is further dependent on the presence of biomolecules *N_x_* within the cytoplasm such as proteins and amino acids where the total osmotic pressure is thus Π = Π*_p_* + Π*_x_*. As the osmotic pressure evolves, it promotes a water flux across the membrane, described by *J_v_* = −*L_p_*_,*m*_(Δ*P* − ΔΠ), where *L_p_*_,*m*_ is the water permeability of the cell membrane associated with aquaporins, and Δ*P* and ΔΠ are the hydrostatic and osmotic pressure differences across the cell membrane, respectively. As water enters a cell, the increase in fluid volume stretches the cell membrane. We treat the membrane and cortex as a single mechanical structure. The constitutive law of the cortical structure can be written as *σ* = *σ_p_* + *σ_a_*, where *σ_a_* is the active stress associated with myosin contractility and *σ_p_* is the passive stress predominantly associated with deformation of the actin network expressed as 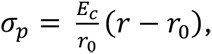 where *E_c_* is the effective stiffness and *r* is the cell radius. From Laplace’s law, the hydrostatic pressure is related to this cortical stress through the relation Δ*P* = 2*σ*ℎ/*r*, where *r* is the cell radius and ℎ is the cortical thickness. In a spherical cell, the change in cell volume follows as *V̇* = *AJ_v_*, where *A* is the surface area of the cell, leading to:

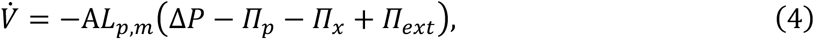

where Π*_ext_* is the osmotic pressure of ions in the microenvironment. The transport of these ions across the membrane is controlled through several active and passive biomechanisms that actively coordinate to maintain intracellular osmolarity. Building on our previous work ^56^, we consider electrochemical fluxes that govern such transport arising from the classic pump-leak model with the form:

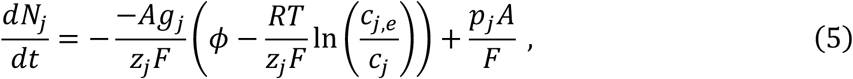

Here, molar fluxes are defined for dominant ions associated with cellular volume control, *φ* is the membrane potential, *A* is the cell surface area, *F* is the Faraday constant, *z*_j_ is the associated ion valence, *p*_j_ is the strength of the pump current, and *g*_j_ = *g*_j0_ + *g*_j,*k*_ are the ion conductances for a given species. The log-based term is a form of the Nernst equation, determining the electrical potential of the species as a function of internal and external (*e*) concentrations. In this framework, we consider the influence of mechanically regulated channels on volume regulation. Piezo1 opens under tension to permit cation transport with an electrochemical gradient, with a mechanosensitive factor 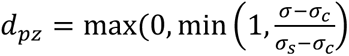 increasing between a threshold (*σ_c_* ) and saturation (*σ_s_* ) stress before plateauing. Piezo1 activation *a_pz_* is transient to prevent cell cytotoxicity, with a rate described by:

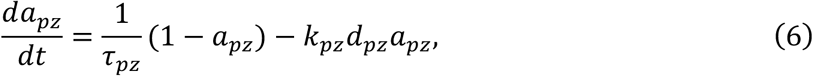

where *k_pz_* governs the rate of suppression after stimulation. The net channel conductance follows as *g_j_*_,*pz*_ = *β_j_*_,*pz*_*α_pz_* where *β_j_*_,*pz*_ is the max channel conductance and *α_pz_* = *a_pz_d_pz_* is the opening probability. Activation of the VRAC is similarly described, with a threshold-shifted Hill function for the mechanosensitive factor given as 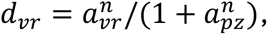 where 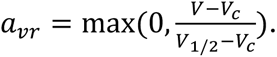 The change in VRAC opening probability then follows as

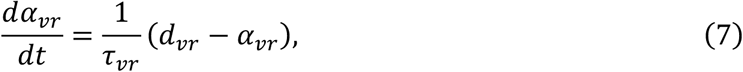

and the associated channel conductance is *g_j_*_,*vr*_ = *β_j_*_,*vr*_*α_vr_*. VRAC also facilitates the transport of neutral osmolytes *N_n_* such as taurine ^65^ whose depletion contributes to volume regulation. Transport of these osmolytes is described through an additional flux

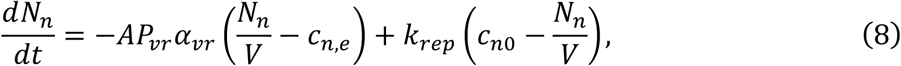

where *P_vr_* is the VRAC permeability to these species and *k_rep_* determines the rate at which they are replenished in the cell to a characteristic concentration *c_n_*_0_. Finally, Mori (2011)^83^ demonstrated that the membrane potential may be approximated by

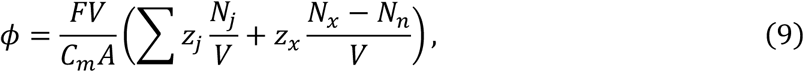

where *C_m_* is the membrane capacitance per unit area. *Model Parameters:* We confine associated modelling parameters to ranges reported by McEvoy et al ^56^. We assume the external ion concentrations remain at physiological levels, such that 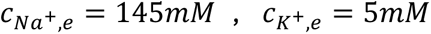 and 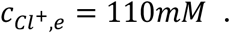 The conductivity coefficients are *g_Na_*_,0_ = 0.1 *S*/*m*^2^, *g_Cl_*_,0_ = 2 *S*/*m*^2^, and *g_K_*_,0_ = 0.45 *S*/*m*^2^, with *C_m_* = 0.01 *F*/*m*^2^. The pump currents reflect the 3:2 stoichiometry of *Na* − *K* ATPase with *p_Na_* = −12.5 *mA*/*m*^2^ and *p_K_* = 8.3 *mA*/*m*^2^. The number of charged non-permeable biomolecules is (*N_x_* − *N_n_*) = 4.75x 10^−14^ *mol* with a mean valence *z_x_* = −1.5, and the reference number of neutral permeable osmolytes is *N_n_* = 3x 10^−14^ *mol*. The reference membrane water permeability due to aquaporins L_p,ref_ = 1×10^−13^ m.s^-1^Pa^-1^. The max conductance of the piezo1 channels *β*_j,*Na*_ = 2 *S*/*m*^2^ with associated parameters σ_c_ = 1.125 kPa, σ_s_ = 1.85 kPa, *τ_pz_* = 20 *s*, and *k_pz_* = 0.45 *s*^−1^. The max permeability and conductance of VRAC are *P_vr_*_,*ref*_ = 6.54 x 10^−7^ *m*/*s* and *β_Cl_*_,*vr*_ = 0.1 *S*/*m*^2^ with associated parameters *V_c_* = 625 µm^3^, *V*_1/2_ = 665 µm^3^, *n* = 4, and *τ_vr_* = 400 *s*. For the timescales investigated in this study, we neglect osmolyte replenishment with *k_rep_* = 0 *s*^−1^. The reference cell volume is 460 µm^3^, with a cell cortex thickness given as 0.6 µm and a stiffness of 2.5 *kPa*. The initial active tension *σ_a_* = 300 *Pa*, increasing to 1 *kPa* at the onset of a stimulus. As intracellular calcium concentrations are too low to materially affect osmotic pressure, we do not directly analyse them in this framework, but the model can readily be extended to evaluate secondary signalling arising from calcium transients.

### Statistical analysis

Statistical analysis was performed using Prism software version 10.6.1 (GraphPad). The variance between the mean values of two groups was evaluated using the unpaired Student’s *t*-test. For assessment of more than two groups, we used one-way analysis of variance (ANOVA) test. A *P* value of <0.05 was considered statistically significant. Statistic details can be found in each figure legend and corresponding Source Data file. Quantitative image analyses of lateral plate mesoderm nuclear tracking and of calcium imaging were performed in Python 3.10.19 with SciPy and statsmodels, as described in the corresponding Methods sections; these used Welch’s t-test (unequal variances) and, where stated, Holm Bonferroni correction or Type II two-way ANOVA with Tukey’s HSD.

## Data Availability

The data supporting the finding of this study are available within the article and Supplementary Information or from the corresponding author upon reasonable request. Source data are provided with this paper.

## Code availability

The image-analysis pipelines used in this study, including nuclear tracking of lateral plate mesoderm cells and calcium imaging analysis, are available at https://github.com/yanc0913/AQP_HSPC and archived at Zenodo https://doi.org/10.5281/zenodo.22639007. The repository documents every analysis parameter, the segmentation, tracking, midline-detection and event-detection methods, and the statistical tests, and includes the Fiji macro used for calcium ROI measurement.

## Acknowledgements

We thank members of the Phng Lab for discussions and suggestions; Emi Taniguchi and RIKEN BDR Research Aquarium for technical assistance; Hiroyuki Nakajima and Naoki Mochizuki for providing the *Tg(fli1:GAL4FF);Tg(UAS:GCaMP7a)* zebrafish line; Anne Schmidt, Shigetomo Fukuhara, Kiryl Piatkevich, Massimo Santoro and Yasushi Okada for sharing plasmids; and Leonard Zon, Chloé Baron, Nathan Lawson and Fumio Motegi for critical feedback. This work was supported by intramural funding from RIKEN BDR (to L-K.P.), JSPS Grants-in-Aid for Scientific Research grants (22H022624 and 22H05168 to L-K.P; 25K09658 to I.K.) and ERC Starting Grant (101116234 to E.M.). Fig. 6h and Fig. 7i were created in BioRender. Phng, L. (2027) https://BioRender.com/d8nr3x1.

## Contributions

L.K.P. and I.K conceptualised the project. I.K., Y.C., R.K., J.D.S., H.W. and G.C. performed experiments. E.M. conducted computational simulations, analyses and prepared the corresponding figures and text. Y.C. developed Fiji and Python macros for image processing and analyses. K.K. provided reagent. I.K., Y.C. and L.K.P. wrote the manuscript. L.K.P. acquired funding and supervised the project.

