## Supplemental Figures for "Osmo-hydraulic volume regulation ensures robust endothelial-to-haematopoietic transition"

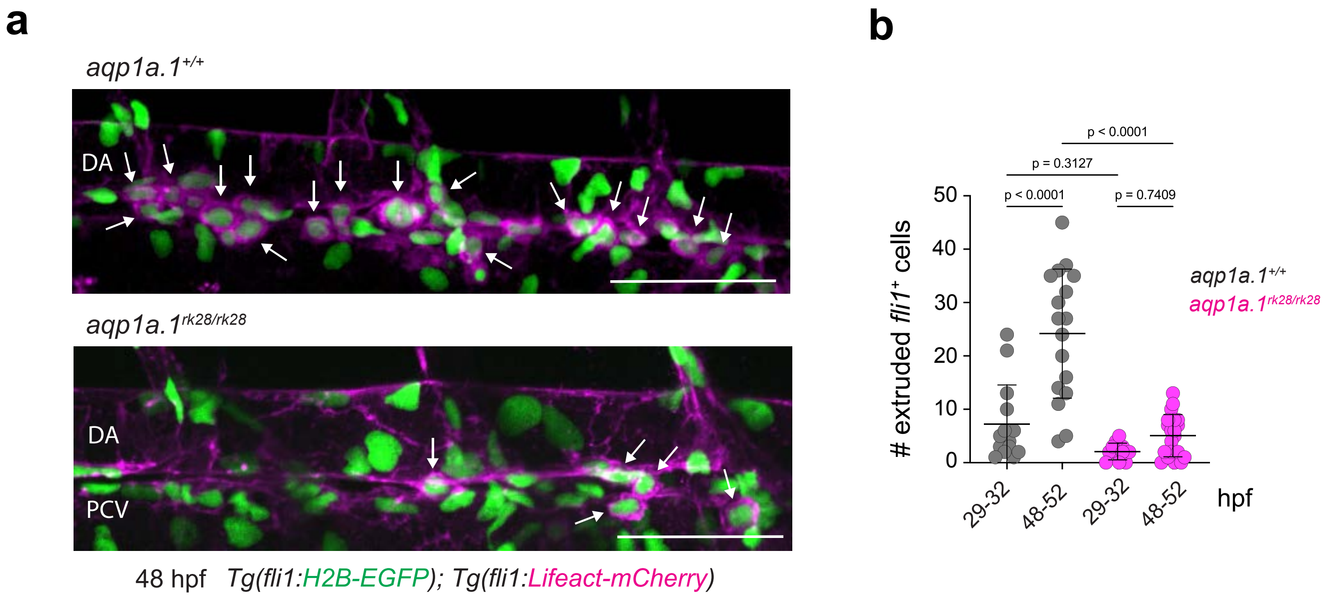


**Supplementary Fig. 1. The number of *fli1*$^{+}$ cells extruded from the DA is decreased in *aqp1a.1*$^{rk28}$ mutant.** **a** Representative maximum intensity projection images of *aqp1a.1*$^{+/+}$ and *aqp1a.1*$^{rk28/rk28}$ embryos in *Tg(fli1:H2B-EGFP);Tg(fli1:Lifeact-mCherry)* background at 48 hpf. White arrows indicate extruded *fli1*$^{+}$ cells. DA, dorsal aorta; PCV, posterior cardinal vein. Scale bars, 50 μm. **b** Quantification of extruded *fli1*$^{+}$ cells in *aqp1a.1*$^{+/+}$ and *aqp1a.1*$^{rk28/rk28}$ embryos at 29-32 hpf (*aqp1a.1*$^{+/+}$, n=14 embryos; *aqp1a.1*$^{rk28/rk28}$, n=12 embryos) and 48-52 hpf (*aqp1a.1*$^{+/+}$, n=17 embryos; *aqp1a.1*$^{rk28/rk28}$, n=20 embryos). Data were collected from 2 independent experiments and presented as mean ± SD. Statistical significance was determined by ordinary one-way ANOVA with Sidak's multiple comparisons test.

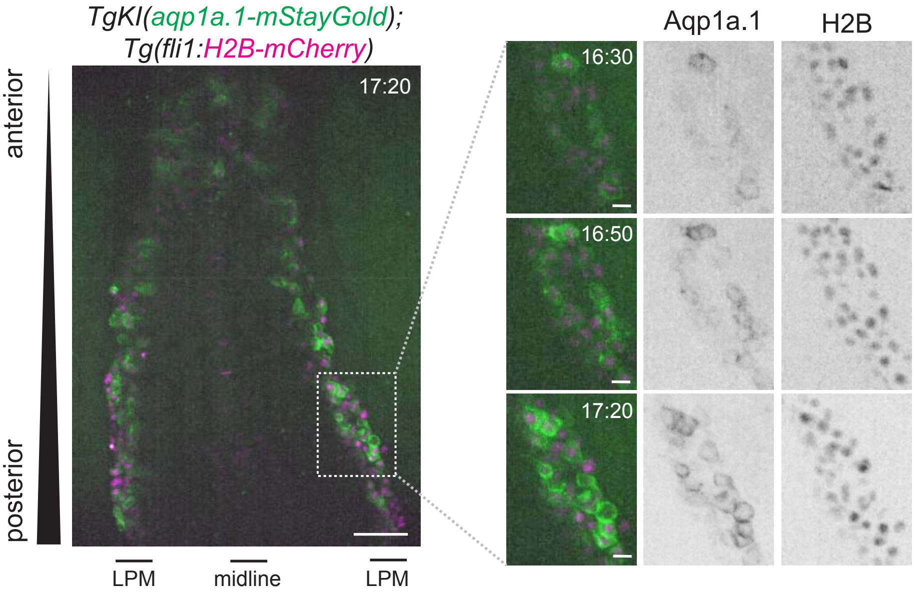


**Supplementary Fig. 2. Aqp1a.1 protein is expressed in LPM-derived *fli1*⁺ cells.** Aqp1a.1 expression in the plasma membrane increases in migrating fli1+ cells derived from the posterior LPM. A magnified view of the boxed region is shown to the right as still images extracted from time-lapse movie of *TgKI(aqp1a.1-mStayGold);Tg(fli1:H2B-mCherry)* embryo. Movie was taken from 13 to 17 hpf. Time, hours:minutes. LPM, lateral plate mesoderm. Scale bars, 10 μm (dashed box) and 50 μm.

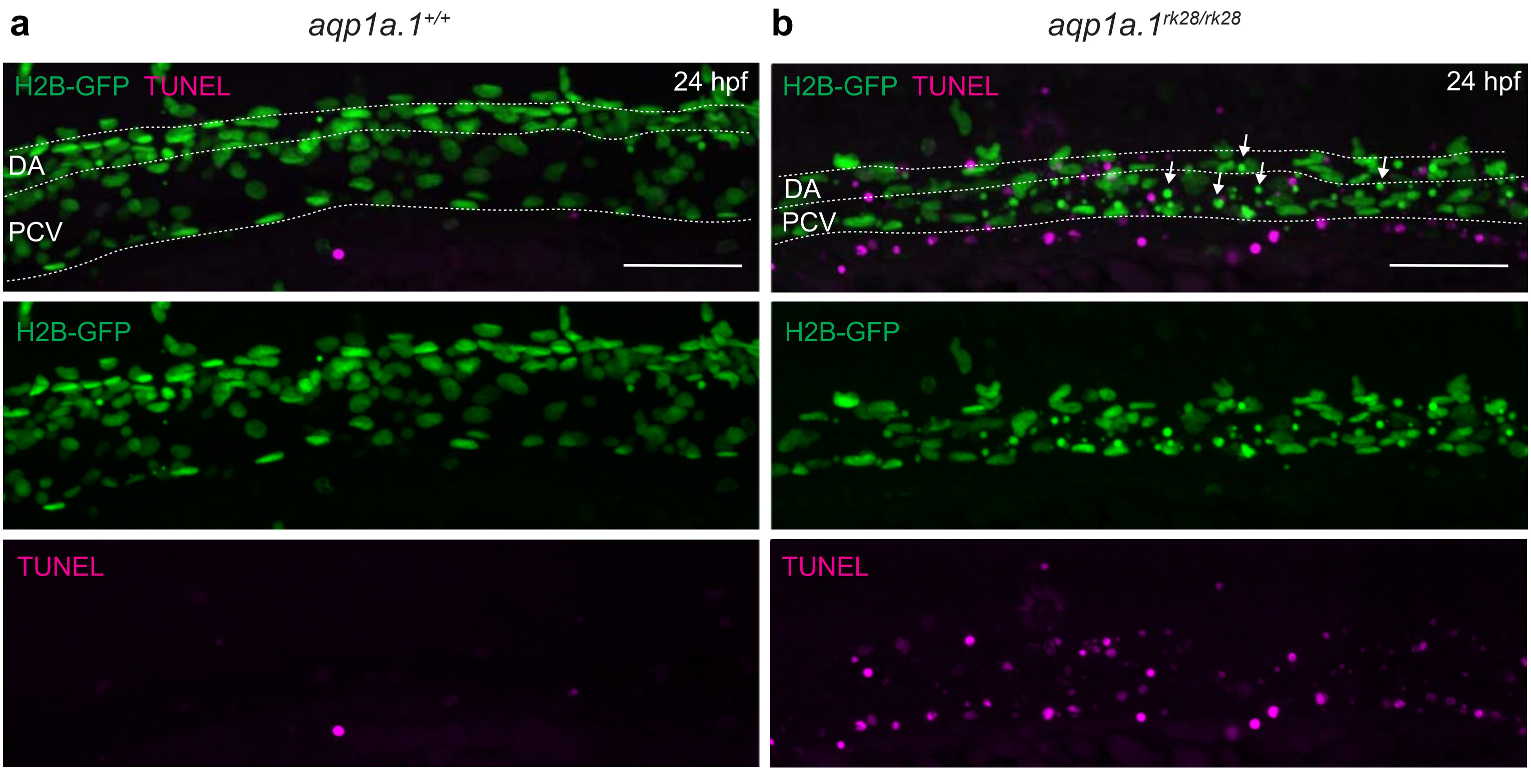


**Supplementary Fig. 3. Cell death in the axial vessels of aqp1a.1rk28/rk28 embryos.** TUNEL (magenta) and anti-GFP immunofluorescent staining in *aqp1a.1$^{+/+}$;Tg(fli1:H2B-EGFP)* (**a**) and *aqp1a.1$^{rk28/rk28}$;Tg(fli1:H2B-EGFP)* (**b**) embryos at 24 hpf. In *aqp1a.1* mutants, fragmented GFP$^{+}$ endothelial cell nuclei (white arrows) at the DA and PCV are not always stained for TUNEL (magenta, apoptotic cells). Region of the DA between ISVs no. 6 and 11 is shown. DA, dorsal aorta; PCV, posterior cardinal vein. Dotted lines mark boundaries of the DA and PCV. Scale bar, 50 μm.

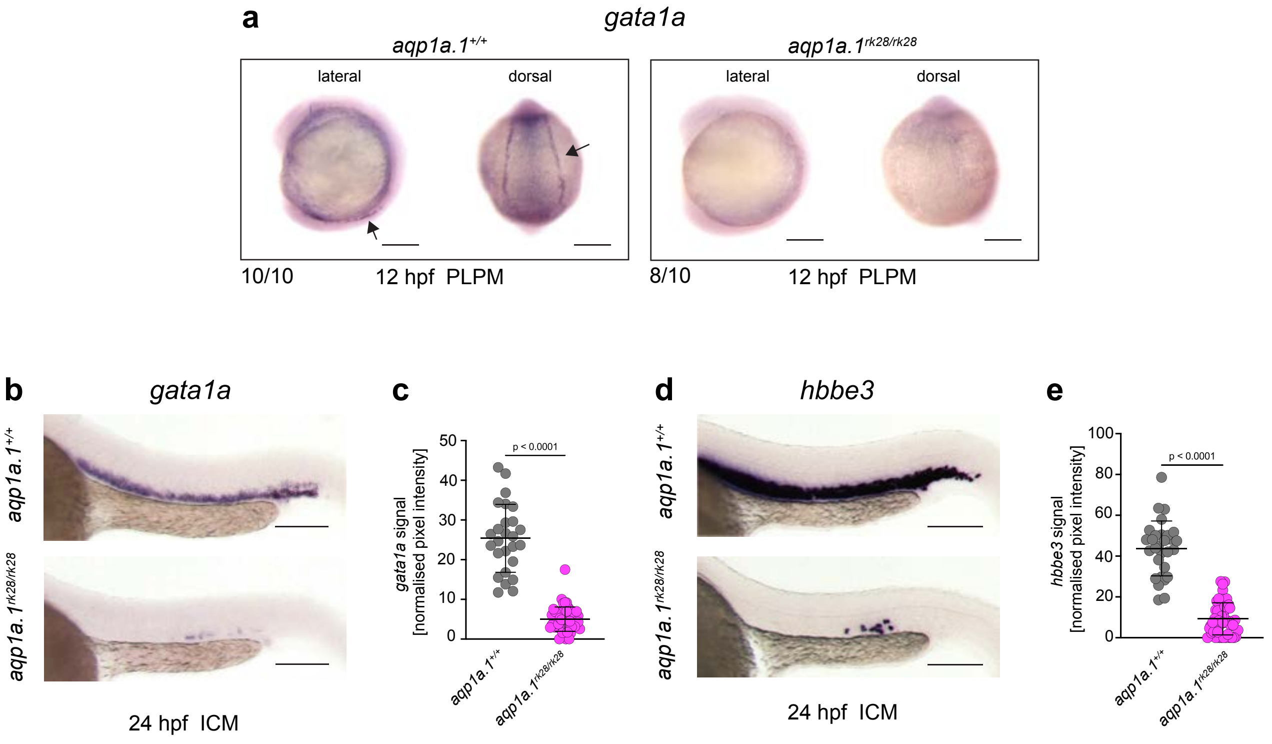


**Supplementary Fig. 4. *aqp1a.1$^{rk28/rk28}$* embryos show reduced expression of erythroid markers.** **a** Expression of *gata1a* mRNA in *aqp1a.1$^{+/+}$* and *aqp1a.1$^{rk28/rk28}$* embryos at 12 hpf. Black arrows denote *gata1a* expression in PLPM. **b** Expression of *gata1a* mRNA in *aqp1a.1$^{+/+}$* and *aqp1a.1$^{rk28/rk28}$* embryos in ICM at 24 hpf. **c** Quantification of expression of *gata1a* in *aqp1a.1$^{+/+}$* (n=25 embryos) and *aqp1a.1$^{rk28/rk28}$* (n=50 embryos) embryos at 24 hpf. **d** Expression of *hbbe3* mRNA in *aqp1a.1$^{+/+}$* and *aqp1a.1$^{rk28/28}$* in ICM at 24 hpf. **e** Quantification of *hbbe3* mRNA expression in *aqp1a.1$^{+/+}$* (n=29 embryos) and *aqp1a.1$^{rk28/28}$* (n=48 embryos) embryos in ICM at 24 hpf. Data are collected from two (**e**) or three (**c**) independent experiments and presented as mean ± SD. Statistical significance was determined by two-tailed unpaired t-test. PLPM, posterior lateral plate mesoderm; ICM, intermediate cell mass. Scale bars, 200 μm (**a**) and 150 μm (**b**, **d**).

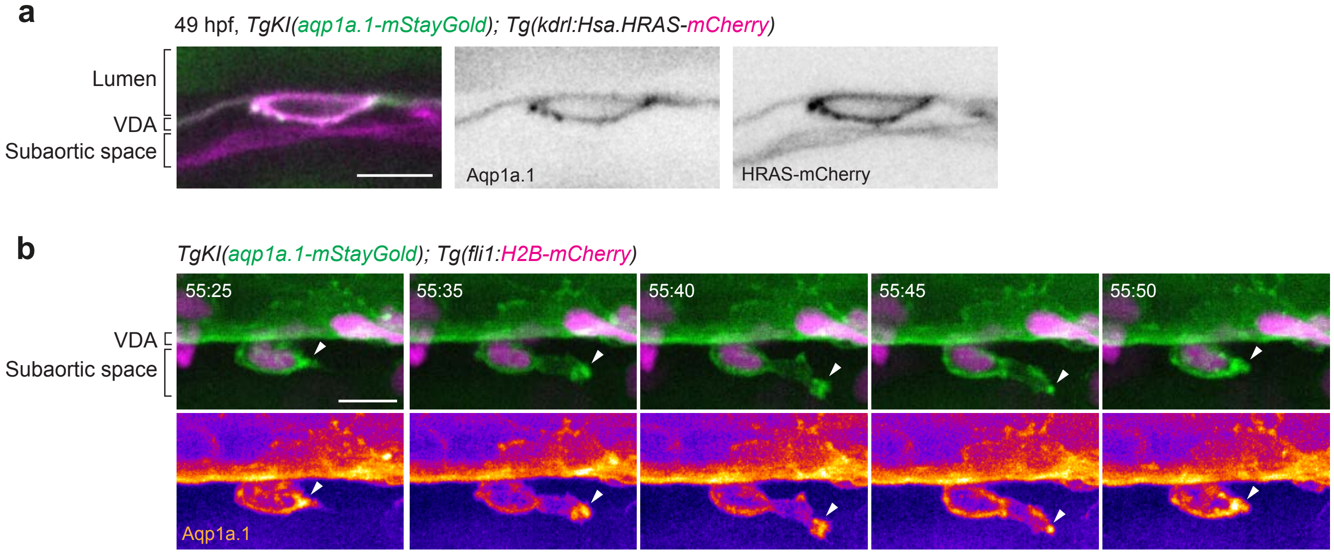


**Supplementary Fig. 5. Aqp1a.1 is localised at the plasma membrane of HEC and HSPC. a** Aqp1a.1 protein is localised to the plasma membrane of HEC undergoing EHT. Representative maximum intensity projection image of HEC in *TgKI(aqp1a.1:mStayGold);Tg(kdrl:Hsa.HRAS-mCherry)* embryo at 49 hpf. **b** Aqp1a.1 protein is localised to the plasma membrane of nascent HSPC in the subaortic space. Still images from time-lapse movie of *TgKI(aqp1a.1-mStayGold);Tg(fli1:H2B-mCherry)* embryo. Movie was taken from 53 to 61 hpf. Arrowhead, Aqp1a.1 enrichment at leading edge of cell protrusion. Time, hour:minutes. HEC, haemogenic endothelial cell; HSPC, haematopoietic stem and progenitor cell; VDA, ventral floor of the dorsal aorta. Scale bars, 10 μm.

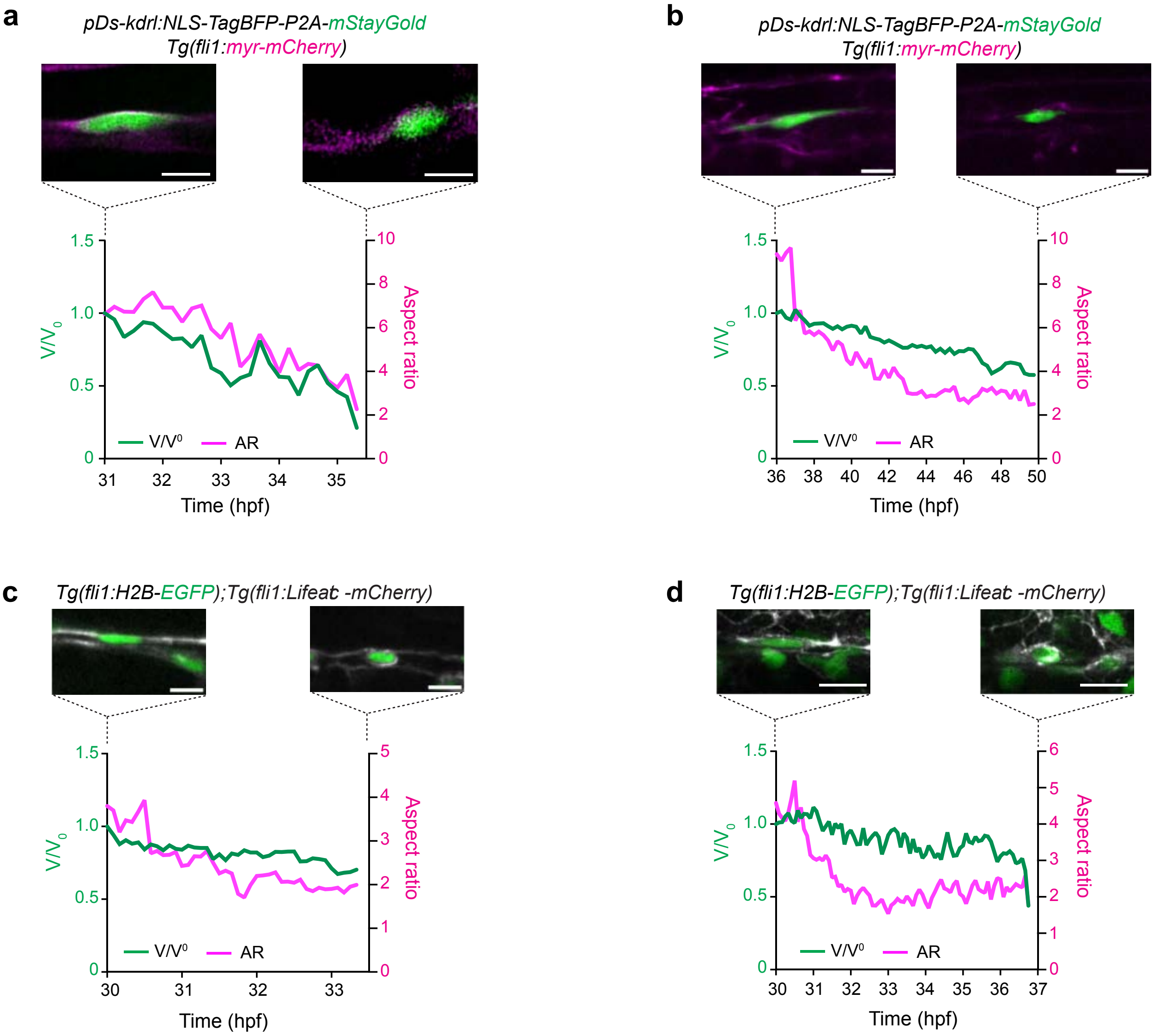


**Supplementary Fig. 6. Haemogenic endothelial cell (HEC) volume decreases during EHT. a-b** Graphs show changes in cell volume and cell aspect ratio of individual HECs from 31 to 35 hpf (**a**) and 36 to 50 hpf (**b**). HEC was transiently labelled by injection of *pDs-kdrl:NLS-TagBFP-P2A-mStayGold* plasmid into *Tg(fli1:myr-mCherry)* embryos. Deconvolved images (single z-plane) of labelled HEC (**a**) and maximum intensity projection (**b**) are shown. **c-d** Graphs show changes in nuclear volume and nuclear aspect ratio of individual HECs in the DA of *Tg(fli1:H2B-EGFP);Tg(fli1:Lifeact-mCherry)* between 30 to 33.5 hpf (**c**) and between 30 to 37 hpf (**d**). Maximum intensity projections are shown. Scale bars, 10 μm.

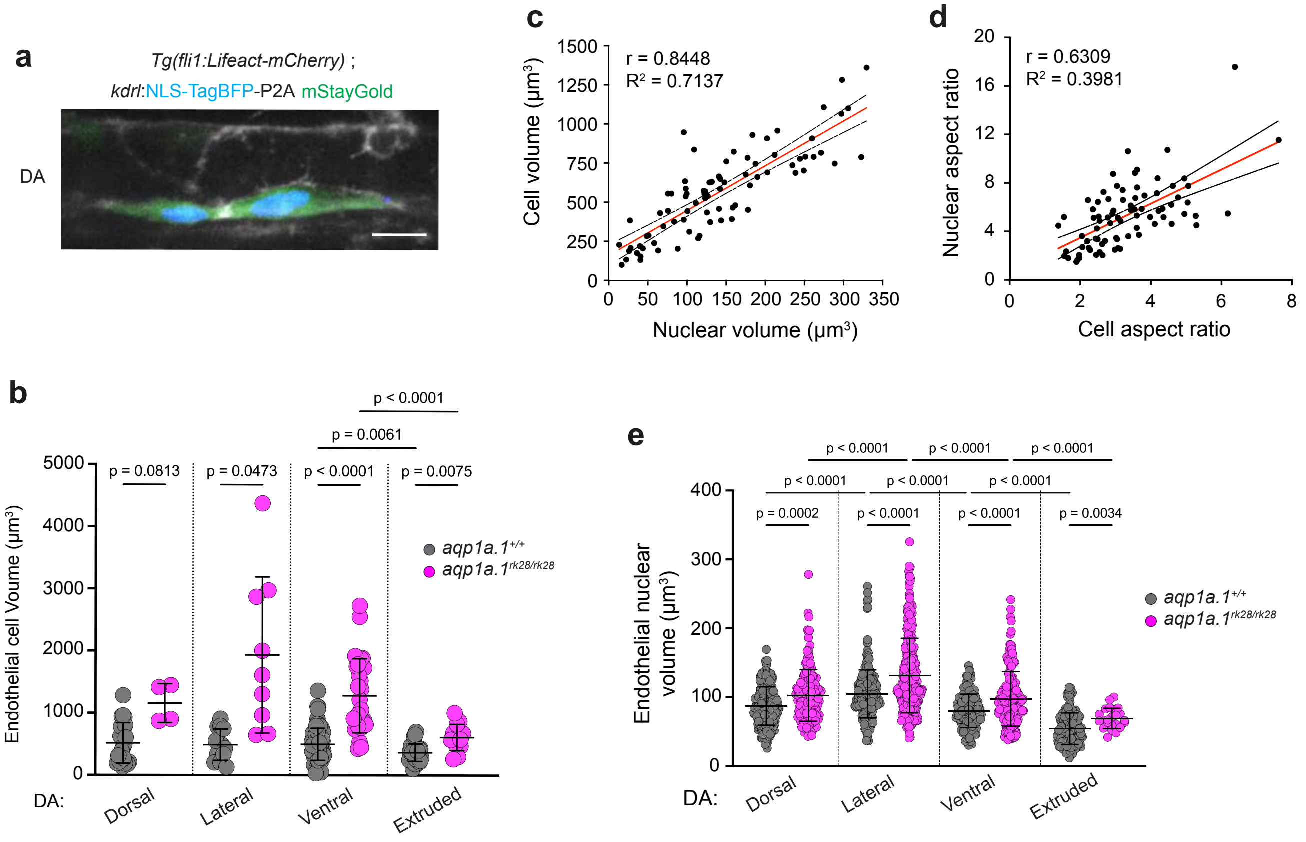


**Supplementary Fig. 7. Aqp1a.1 deficiency triggers EC swelling in the dorsal aorta. a** Nucleus and cytoplasm of single cell were dual labelled by injection of *kdrl:NLS-TagBFP-P2A-mStayGold* plasmid into *Tg(fli1:Lifeact-mCherry)* embryos at 30 hpf, allowing segmentation of nucleus and cell body, respectively. DA, dorsal aorta. Scale bar, 10 µm. **b** Quantification of endothelial cell volume in $aqp1a.1^{+/+}$ and $aqp1a.1^{rk28/rk28}$ embryos at 30-33 hpf ($aqp1a.1^{+/+}$: n=23 dorsal, 13 lateral, 79 ventral and 29 extruded cells from 38 embryos; $aqp1a.1^{rk28/rk28}$: n=4 dorsal, 9 lateral, 29 ventral and 13 extruded cells from 28 embryos). ECs were labelled by transient injection of either *kdrl:NLS-TagBFP-P2A-mStayGold* or *Runx1+23enh:mStayGold* plasmids into $aqp1a.1^{+/+}$ or $aqp1a.1^{rk28/rk28}$ embryos in *Tg(fli1:Lifeact-mCherry)* background. Data were collected from 7 ($aqp1a.1^{+/+}$ embryos) and 11 ($aqp1a.1^{rk28/rk28}$ embryos) independent experiments and presented as mean ± SD. Statistical significance was determined by Brown-Forsythe and Welch ANOVA with Dunnett's T3 multiple comparisons test. **c**-**d** Linear correlation between nuclear and cell volume (**c**), and nuclear and cell aspect ratio (**d**) of ECs of the DA at 30-33 hpf (n=80 cells from 18 embryos, 7 independent experiments). **e** Quantification of nuclear volume of ECs in $aqp1a.1^{+/+}$ and $aqp1a.1^{rk28/rk28}$ embryos in *Tg(fli1:H2B-EGFP);Tg(fli1:Lifeact-mCherry)* background at 48-52 hpf ($aqp1a.1^{+/+}$: n=266 dorsal, 238 lateral, 164 ventral and 165 extruded cells from 17 embryos; $aqp1a.1^{rk28/rk28}$: n=169 dorsal, 320 lateral, 173 ventral and 23 extruded cells from 24 embryos). Data were collected from 2 independent experiments and presented as mean ± SD. Statistical significance was determined by Brown-Forsythe and Welch ANOVA with Games-Howell multiple comparisons test.

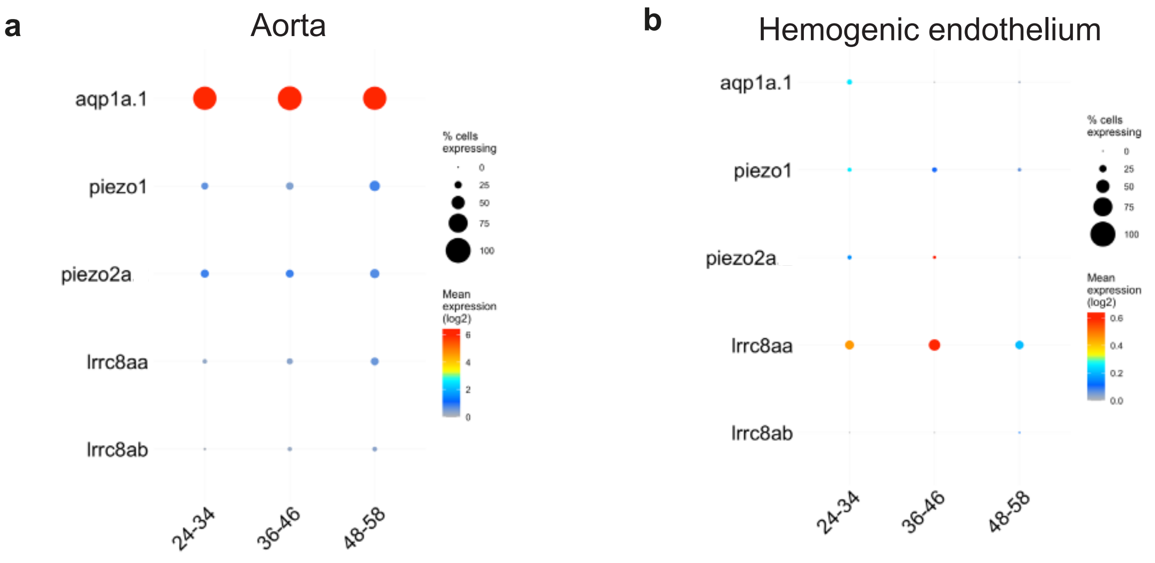


**Supplementary Fig. 8. Expression of ion channels in zebrafish aorta and hemogenic endothelium.** Expression of *aqp1a.1*, *piezo1*, *piezo2a*, *lrrc8aa* and *lrrc8ab* in aorta (hema.18) (**a**) and haemogenic endothelium (hema.21) (**b**) of wild-type zebrafish embryos at 24-58 hpf. Single-cell gene expression data were collected from Daniocell (https://daniocell.nichd.nih.gov).

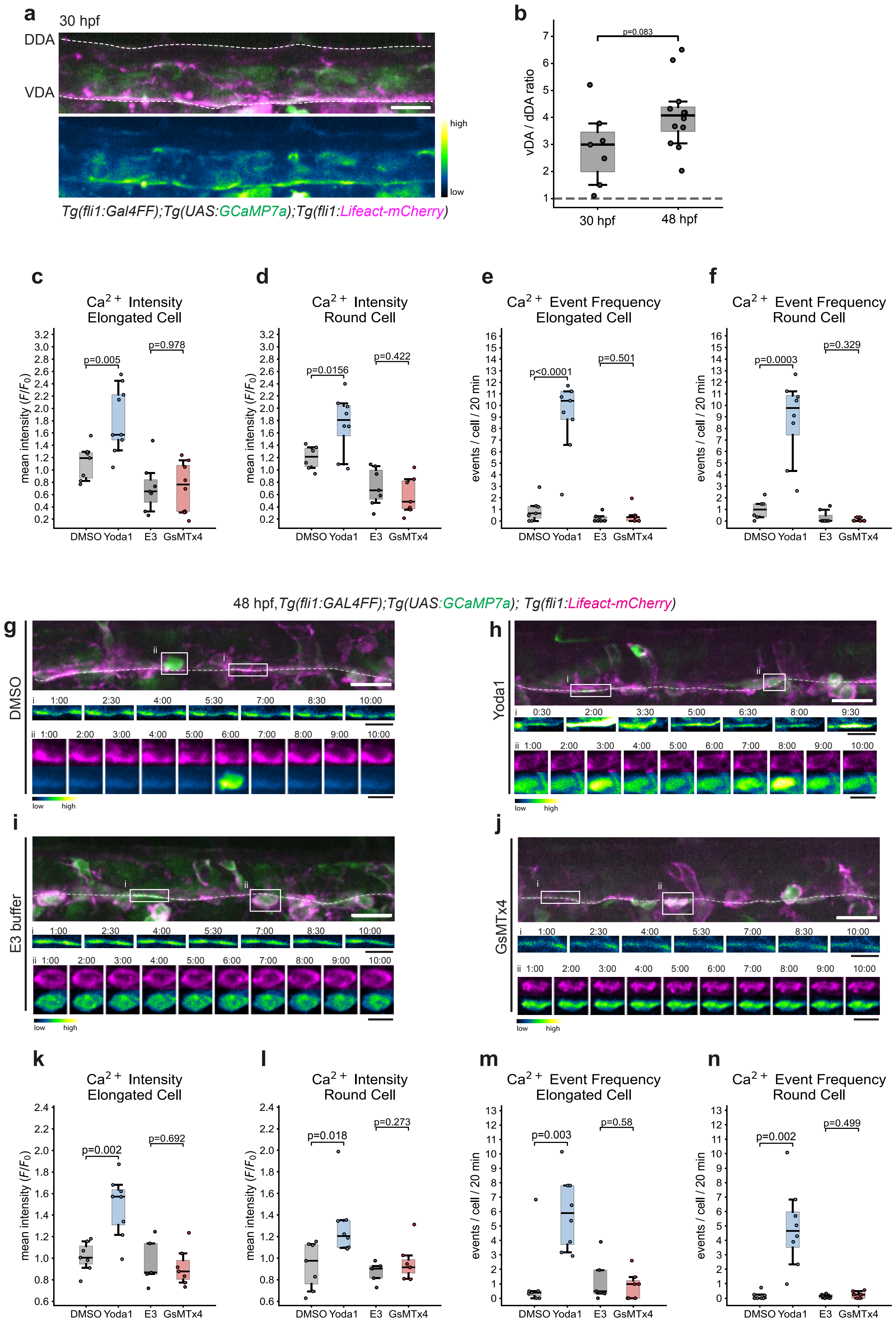

a
30 hpf
DDA
VDA
high
low
Tg(fli1:Gal4FF);Tg(UAS:GCaMP7a);Tg(fli1:Lifeact-mCherry)
b
p=0.083
vDA / dDA ratio
30 hpf
48 hpf
c
Ca2+ Intensity Elongated Cell
p=0.005
p=0.978
mean intensity (F/F0)
DMSO Yoda1 E3 GsMTx4
d
Ca2+ Intensity Round Cell
p=0.0156
p=0.422
mean intensity (F/F0)
DMSO Yoda1 E3 GsMTx4
e
Ca2+ Event Frequency Elongated Cell
p<0.0001
p=0.501
events / cell / 20 min
DMSO Yoda1 E3 GsMTx4
f
Ca2+ Event Frequency Round Cell
p=0.0003
p=0.329
events / cell / 20 min
DMSO Yoda1 E3 GsMTx4
48 hpf, Tg(fli1:GAL4FF);Tg(UAS:GCaMP7a); Tg(fli1:Lifeact-mCherry)
g
DMSO
h
Yoda1
i
E3 buffer
j
GsMTx4
low
high
k
Ca2+ Intensity Elongated Cell
p=0.002
p=0.692
mean intensity (F/F0)
DMSO Yoda1 E3 GsMTx4
l
Ca2+ Intensity Round Cell
p=0.018
p=0.273
mean intensity (F/F0)
DMSO Yoda1 E3 GsMTx4
m
Ca2+ Event Frequency Elongated Cell
p=0.003
p=0.58
events / cell / 20 min
DMSO Yoda1 E3 GsMTx4
n
Ca2+ Event Frequency Round Cell
p=0.002
p=0.499
events / cell / 20 min
DMSO Yoda1 E3 GsMTx4

**Supplementary Fig. 9. Regional $Ca^{2+}$ activity in the dorsal aorta and per-group measurements underlying Fig. 5e-h. a** Representative maximum intensity projection of the DA in *Tg(fli1:GAL4FF);Tg(UAS:GCaMP7a);Tg(fli1:Lifeact-mCherry)* embryos, showing GCaMP7a signal in green (upper) or as a pseudocolour heat map (LUT: Green Fire Blue, lower), and Lifeact-mCherry in magenta (upper). Dashed lines indicate the ventral (VDA) and dorsal (DDA) regions of the DA. Scale bar, 20 µm. **b** VDA/DDA GCaMP7a ration in untreated embryos, taken as the median ratio across the 5 min baseline recording, at 30 and 48 hpf. The dashed line marks a ratio of 1 (30 hpf, n=7 embryos; 48 hpf, n=12 embryos). **c**-**f** Per-group $Ca^{2+}$ measurements at 30 hpf, from which the fold change in Fig. 5e-h are derived: mean GCaMP7a intensity, the $F/F_0$ signal averaged over the 20 min post-treatment window (**c**, elongated; **d**, round), and $Ca^{2+}$ event frequency, expressed as events per cell per 20 min (**e**, elongated; **f**, round). Data are from 4 independent experiments (DMSO, n=23/14 elongated/round cells from 7/6 embryos; Yoda1, n=29/20 cells from 9/8 embryos; E3, n=20/15 cells from 7/7 embryos; GsMTx4, 21/16 cells from 8/7 embryos). **g**-**j** Representative maximum intensity projections of the VDA at 48 hpf during the post-treatment phase, after 30 min in 0.7% DMSO (**g**), 20 µM Yoda1 (**h**), E3 buffer (**i**) or 1 µM GsMTx4 (**j**), displayed as in Fig. 5a-d. Times, min:sec from the start of the post-treatment recording. Scale bar, 20 µm; insets, 10 µm. **k**-**n** The same measurements as c-f at 48 hpf from 4 independent experiments (DMSO, n=20/23 elongated/round cells from 7 embryos; Yoda1, n=28/38 cells from 8 embryos; E3, n=15/24 cells from 6 embryos; GsMTx4, 16/29 cells from 7 embryos). Statistical significance was determined by two-sided Welch's *t*-test on per embryo values, each drug against its matched vehicle. Exact *p* values are shown.

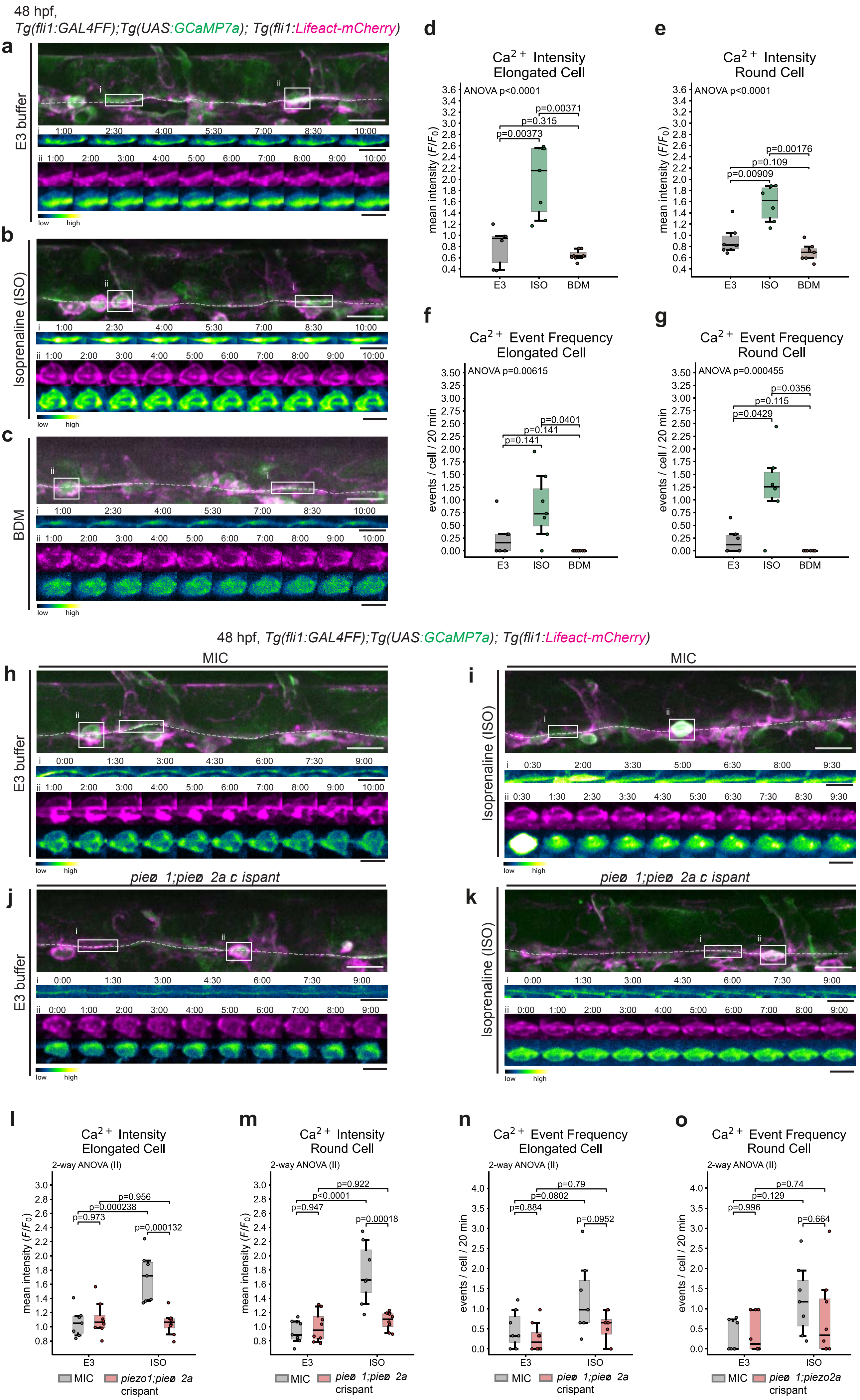

48 hpf,
Tg(fli1:GAL4FF);Tg(UAS:GCaMP7a); Tg(fli1:Lifeact-mCherry)
a
E3 buffer
b
Isoprenaline (ISO)
c
BDM
low high
d
Ca2+ Intensity
Elongated Cell
ANOVA p<0.0001
p=0.00373
p=0.315
p=0.00371
mean intensity (F/F0)
E3 ISO BDM
e
Ca2+ Intensity
Round Cell
ANOVA p<0.0001
p=0.00909
p=0.109
p=0.00176
f
Ca2+ Event Frequency
Elongated Cell
ANOVA p=0.00615
p=0.141
p=0.141
p=0.0401
events / cell / 20 min
g
Ca2+ Event Frequency
Round Cell
ANOVA p=0.000455
p=0.0429
p=0.115
p=0.0356
48 hpf, Tg(fli1:GAL4FF);Tg(UAS:GCaMP7a); Tg(fli1:Lifeact-mCherry)
MIC
h
E3 buffer
i
Isoprenaline (ISO)
piezo1;piezo2a crispant
j
E3 buffer
k
Isoprenaline (ISO)
l
Ca2+ Intensity
Elongated Cell
2-way ANOVA (II)
p=0.973
p=0.000238
p=0.956
p=0.000132
m
Ca2+ Intensity
Round Cell
2-way ANOVA (II)
p=0.947
p<0.0001
p=0.922
p=0.00018
n
Ca2+ Event Frequency
Elongated Cell
2-way ANOVA (II)
p=0.884
p=0.0802
p=0.79
p=0.0952
o
Ca2+ Event Frequency
Round Cell
2-way ANOVA (II)
p=0.996
p=0.129
p=0.74
p=0.664
MIC piezo1;piezo2a crispant

**Supplementary Fig. 10. $Ca^{2+}$ responses to altered cardiac output and per-group measurements underlying Fig. 5i-l. a-c** Representative maximum intensity projections of the VDA in wild-type embryos at 48 hpf during the post-treatment phase after 1 hour in E3 buffer (**a**), 1 hour in 100 µM isoprenaline hydrochloride (ISO, **b**) or 5 min in 50 mM 2,3-butanedione monoxime (BDM, **c**), which arrests the heartbeat, displayed as in Fig. 5a-d. Scale bar, 20 µm; insets, 10 µm. **d-g** Per-group $Ca^{2+}$ measurements in wild-type embryos: mean GCaMP7a intensity (**d**, elongated; **e**, round) and $Ca^{2+}$ event frequency (**f**, elongated; **g**, round). Data are from 4 independent experiments (E3, n=13/17 elongated/round cells from 6 embryos; ISO, 18/22 cells from 7/6 embryos; BDM, 13/31 cells from 7 embryos). Statistical significance was determined by ordinart one-way ANOVA, shown at the top left of each panel, followed by pairwise two-sided Welch's *t*-test with Holm-Bonferroni correction across the three comparisons; adjusted *p* values are shown. **h-k** Representative maximum intensity projections of the VDA at 48 hpf in mock-injected controls (MIC) treated for 1 hour with E3 buffer (**h**) or 100 µM ISO (**i**), and in *piezo1;piezo2a* crispants treated for 1 hour with E3 buffer (**j**) or 100 µM ISO (**k**), displayed as in Fig. 5a-d. Scale bar, 20 µm; insets, 10 µm. **l-o** Per-group $Ca^{2+}$ measurements in mock-injected controls and *piezo1;piezo2a* crispants: mean GCaMP7a intensity (**l**, elongated; **m**, round) and $Ca^{2+}$ event frequency (**n**, elongated; **o**, round). Data are from two independent experiments (MIC-E3, n=23/29 elongated/round cells from 7 embryos; MIC-ISO, n=20/31 cells from 7 embryos; crispant-E3, n=21/36 cells from 8 embryos; crispant-ISO, n=21/28 cells from 8 embryos). Statistical significance was determined by two-way ANOVA (genotype x treatment) followed by Tukey's HSD; adjusted *p* values are shown.

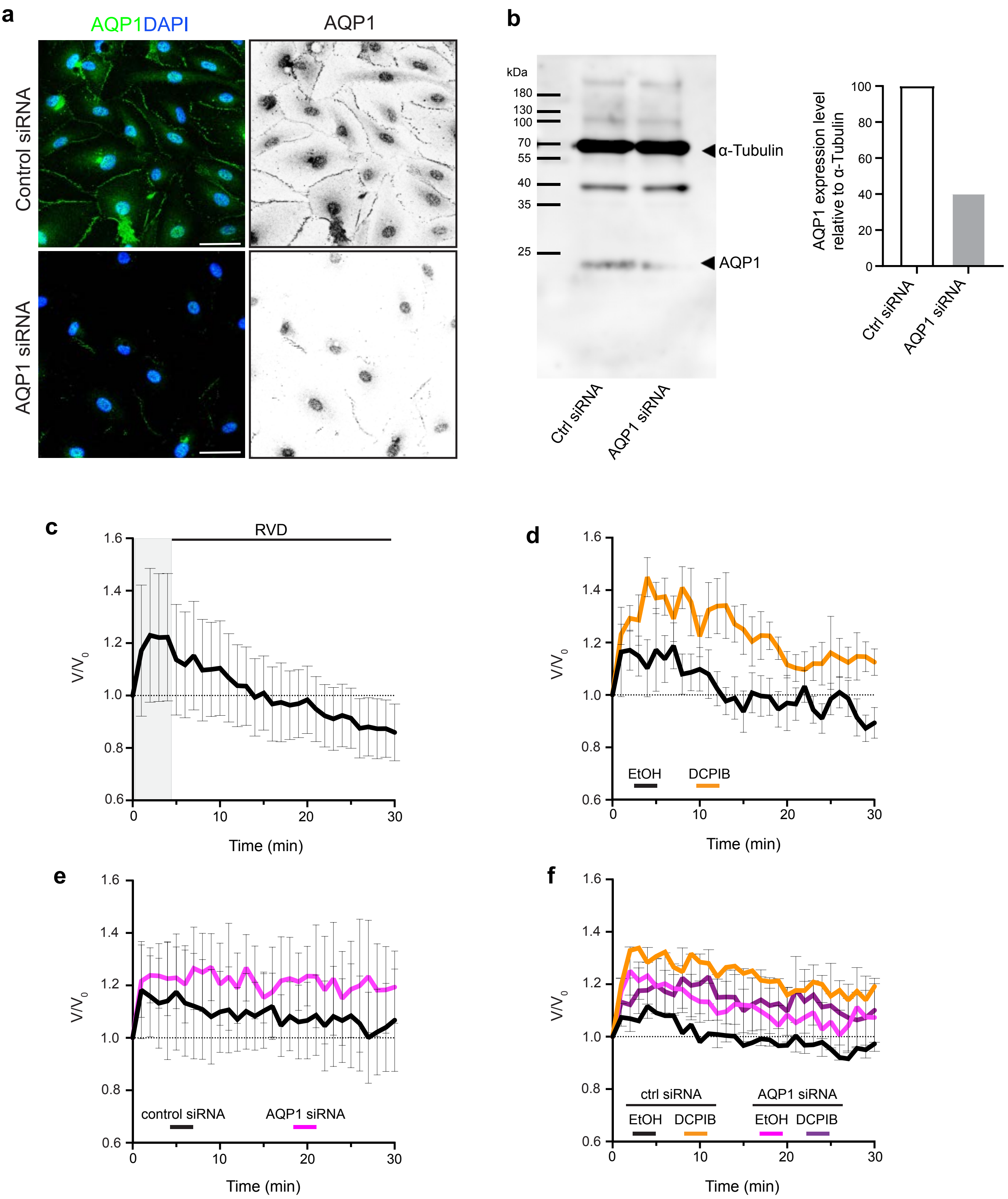
a
AQP1DAPI
AQP1
Control siRNA
AQP1 siRNA
b
kDa
180
130
100
70
55
40
35
25
α-Tubulin
AQP1
Ctrl siRNA
AQP1 siRNA
AQP1 expression level relative to α-Tubulin
100
80
60
40
20
0
Ctrl siRNA
AQP1 siRNA
c
RVD
V/V0
Time (min)
d
EtOH
DCPIB
V/V0
Time (min)
e
control siRNA
AQP1 siRNA
V/V0
Time (min)
f
ctrl siRNA
AQP1 siRNA
EtOH
DCPIB
EtOH
DCPIB
V/V0
Time (min)

**Supplementary Fig. 11. VRAC and AQP1 mediate RVD response of human aortic endothelial cells (HAECs) to a hypotonic shock. a** Representative maximum intensity projection images of HAECs transfected with Control or AQP1 siRNA and stained with anti-AQP1 antibody. Scale bar, 50 µm. **b** Western blot shows a decrease in AQP1 expression level after siRNA knockdown. **c** Effect of hypotonic shock treatment on HAEC nuclear volume. HAECs were exposed to hypotonic solution (water:cell medium, 2:1, v/v) and imaged immediately at 1 min intervals for 30 min (n=24 cells). Gray area highlights initial cell swelling (2-4 min after hypotonic solution addition) followed by RVD (regulatory volume decrease). **d** Effect of hypotonic shock on HAECs with reduced VRAC activity. Cells were treated with 0.05% EtOH (control, n=35 cells) or 5 µM DCPIB (n=20 cells) for 30 min before exposure to hypotonic solution. **e** Effect of hypotonic shock on AQP1-depleted HAECs. Cells were transfected with ON-TARGETplus Non-targeting Pool siRNA (control siRNA) or ON-TARGETplus Human AQP1 siRNA-SMARTpool (AQP1 siRNA) before exposure to hypotonic solution. Control siRNA, n=58 cells; AQP1 siRNA, n=57 cells. **f** Effect of hypotonic shock on HAECs with combined inhibition of VRAC and AQP1 activity. Control siRNA and AQP1 siRNA HAECs were treated with 0.05% EtOH (control siRNA, n=69 cells; AQP1 siRNA, n=62 cells) or 5 µM DCPIB (control siRNA, n=50 cells; AQP1 siRNA, n=75 cells) for 30 min before exposure to hypotonic solution. Data were collected from two independent experiments with two biological replicates per experiment.

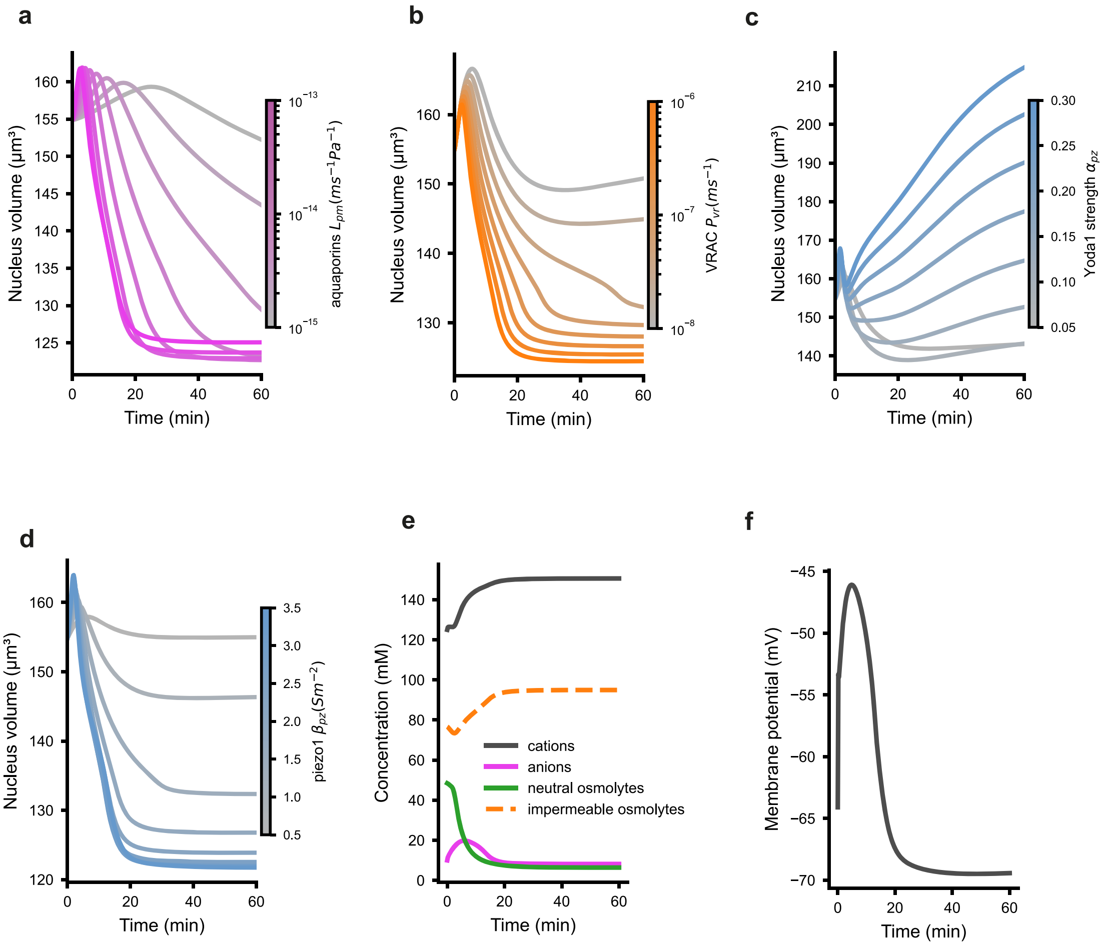


**Supplementary Fig. 12. Parameter studies to analyse the influence of key membrane channels on cell (nucleus) volume. a** Reducing aquaporin-mediated water permeability slows the volume response to osmotic-pressure changes. **b** Reducing Reducing VRAC permeability suppresses osmolyte and anion efflux, limiting volume loss. **c** Sustaining Piezo1 opening with Yoda1 drives elevated cation influx and cell swelling. **d** Reducing Piezo1 expression limits cation influx, which may not drive sufficient swelling to activate VRAC. **e-f** Under control conditions, cation concentration rises with Piezo1 stimulation, while anion and neutral-osmolyte concentrations fall with VRAC activation (**e**); the fixed amount of negatively-charged impermeable osmolyte means its concentration changes with cell volume, in turn influencing membrane potential (**f**).

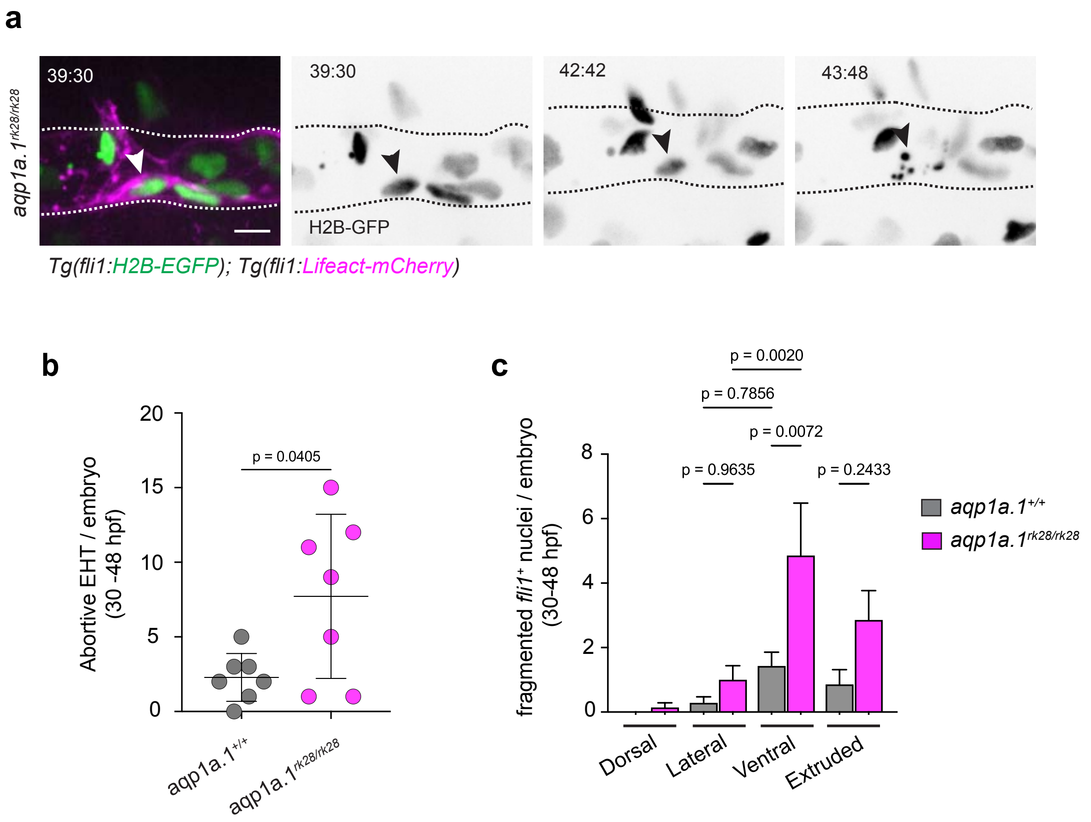


**Supplementary Fig. 13. Reduced water efflux causes endothelial cell death at VDA during EHT.**
**a** Representative maximum intensity projection images from time-lapse movie taken of *aqp1a.1*$^{rk28/rk28}$ embryo in *Tg(fli1:H2B-EGFP);Tg(fli1:Lifeact-mCherry)* background. Time, hours:minutes. Arrowhead, abortive EHT. Serrated lines mark the dorsal aorta boundaries. Scale bar, 10 µm. **b** Quantification of abortive EHT events (nuclear fragmentation) in *aqp1a.1*$^{+/+}$ and *aqp1a.1*$^{rk28/rk28}$ embryos. Data were collected from time-lapse confocal images of embryos from 30 to 48 hpf (*aqp1a.1*$^{+/+}$, n=7 embryos; *aqp1a.1*$^{rk28/rk28}$, n=7 embryos from 4 independent experiments) and presented as mean ± SD. Statistical significance was determined by two-tailed unpaired t-test with Welch's correction. **c** Quantification of endothelial cell death in different regions of the dorsal aorta in *aqp1a.1*$^{+/+}$ (n=7) and *aqp1a.1*$^{rk28/rk28}$ (n=7) *Tg(fli1:H2B-EGFP);Tg(fli1:Lifeact-mCherry)* embryos at 30-48 hpf. Data were collected form time-lapse confocal images of 4 independent experiments and presented as mean ± SEM. Statistical significance was determined by ordinary one-way ANOVA with Sidak's multiple comparisons test.

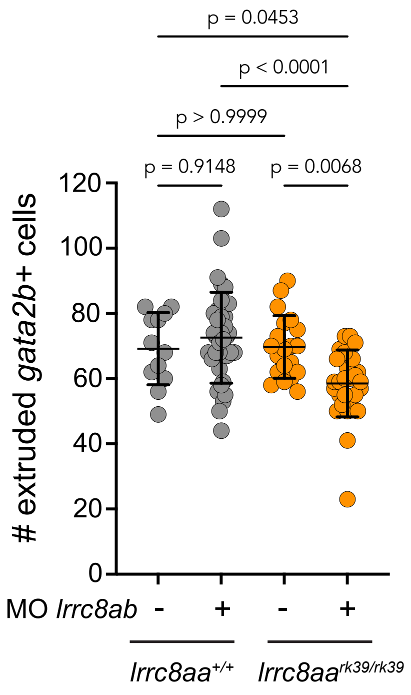


**Supplementary Fig. 14. Depletion of VRAC activity causes a reduction in the number of *gata2b*$^{+}$ cells extruded from the dorsal aorta 48-52 hpf.** Quantification of extruded *gata2b*$^{+}$ cells in *lrrc8aa*$^{+/+}$ and *lrrc8aa*$^{rk39/rk39}$ embryos injected with control or *lrrc8ab* morpholinos (*lrrc8aa*$^{+/+}$; control MO, n=12 embryos; *lrrc8aa*$^{+/+}$*; lrrc8ab* MO, n=36 embryos; *lrrc8aa*$^{rk39/rk39}$; control MO, n=20 embryos; *lrrc8aa*$^{rk39/rk39}$*; lrrc8ab* MO, n=30 embryos). Data were collected from 3 independent experiments and presented as mean ± SD. Statistical significance was determined by ordinary one-way ANOVA with Sidak's multiple comparisons test.

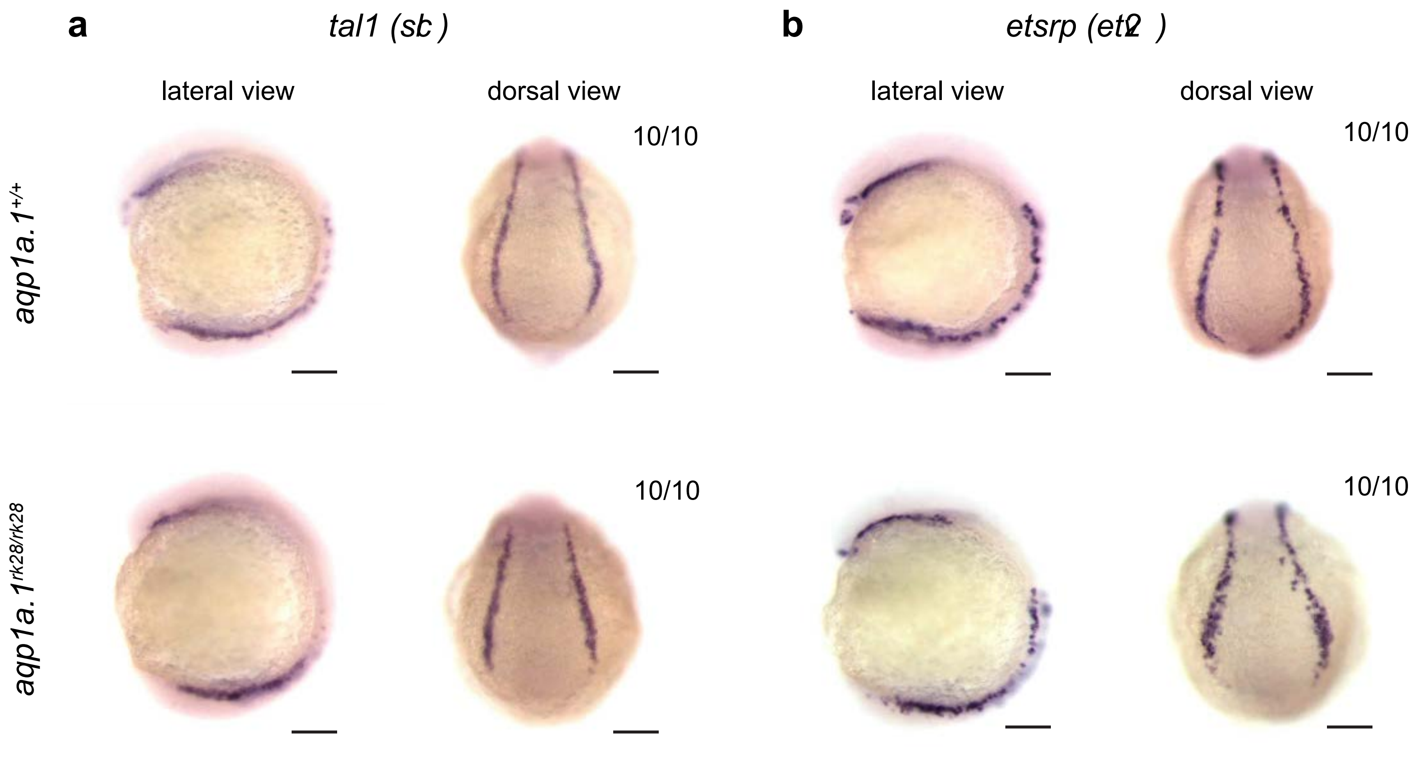


**Supplementary Fig. 15. Expression of *tal1* and *etsrp*.** Expression of tal1 mRNA (**a**) and etsrp mRNA (**b**) in *aqp1a.1*$^{+/+}$ and *aqp1a.1*$^{rk28/rk28}$ embryos at 12 hpf. Scale bar, 150 µm.

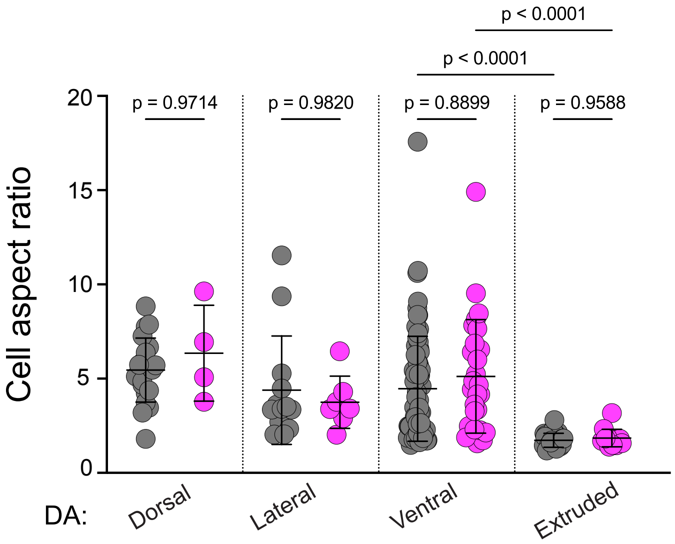


**Supplementary Fig. 16. Cell aspect ratio of endothelial cells in the dorsal aorta at 30-33 hpf.** Single ECs were labelled by injection of *pDs-kdrl:NLS-TagBFP-P2A-mStayGold* plasmid into *Tg(fli1:Lifeact-mCherry)* or *aqp1a.1*$^{rk28/rk28}$;*Tg(fli1:Lifeact-mCherry)* embryos (wild type: n=21 dorsal, 13 lateral, 79 ventral and 29 extruded cells from 38 embryos; *aqp1a.1*$^{rk28/rk28}$: n=4 dorsal, 9 lateral, 28 ventral and 13 extruded cells from 28 embryos). Data are collected from 7 (*aqp1a.1*$^{+/+}$ embryos) and 11 (*aqp1a.1*$^{rk28/rk28}$ embryos) independent experiments and presented as mean ± SD. Statistical significance was determined by Brown-Forsyth and Welch ANOVA with Dunnett's T3 multiple comparisons test.

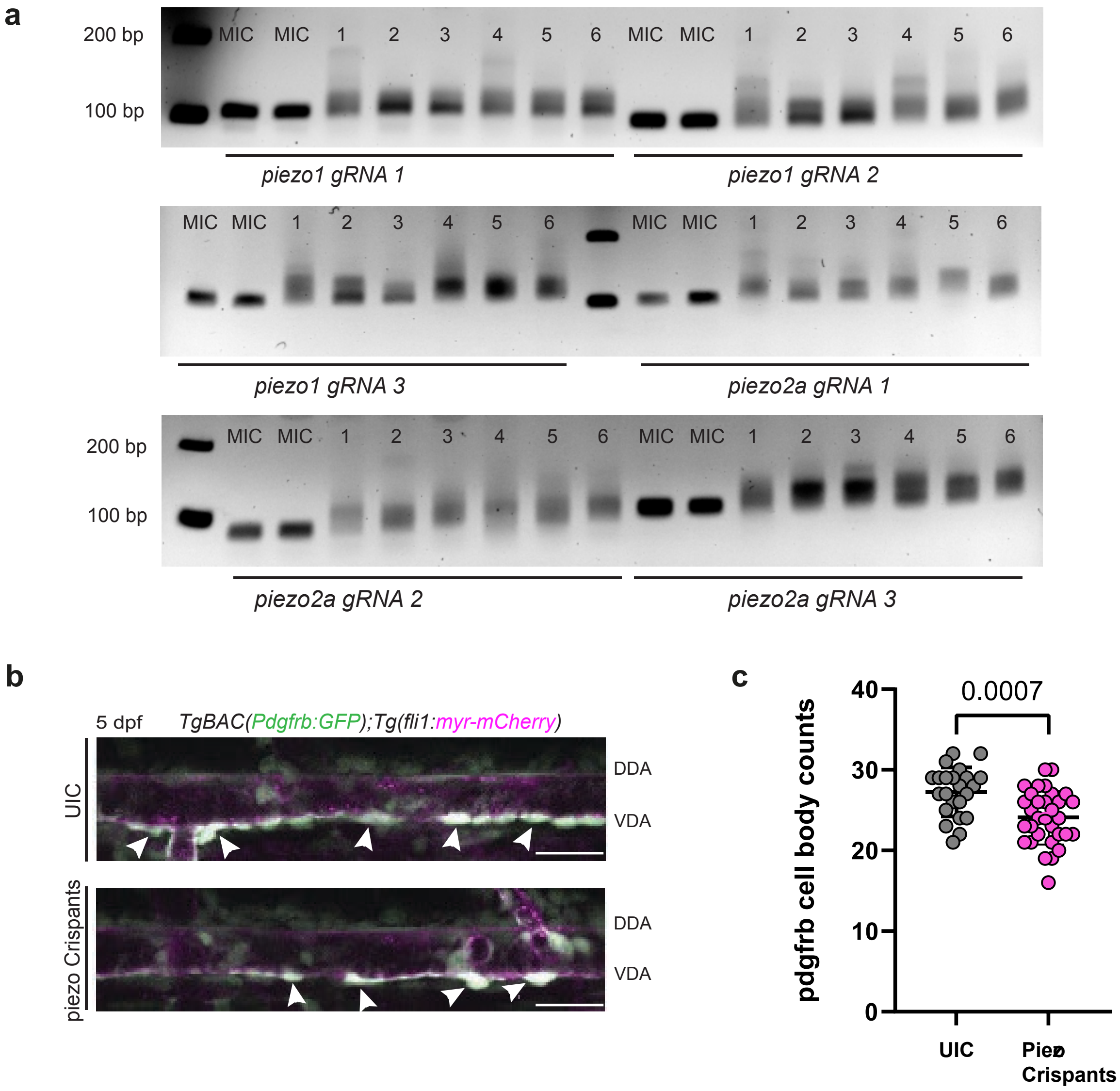


**Supplementary Fig. 17. Evaluation of cutting efficiency at the *piezo1* and *piezo2a* loci. a** PCR screening confirming CRISPR/Cas9-mediated cutting efficiency at the *piezo1* and *piezo2a* loci, using three guide RNAs per gene (six PCR reactions per embryo). UIC, uninjected control, showing a single intact band; lanes 1–6 correspond to individual injected embryos, showing smeared/heteroduplex banding patterns indicative of successful genomic cutting relative to the tight, uncut band observed in the UIC. **b** Representative confocal images of *TgBAC(pdgfrb:EGFP);Tg(fli1:myr-mCherry)* embryos at 5 days post-fertilization (dpf), comparing uninjected controls to *piezo1;piezo2a* crispants. Arrows indicate Pdgfrb-positive mural cells associated with the ventral wall of the dorsal aorta. Scale bar, 20 µm. **c** Quantification of the number of Pdgfrb-positive cells at the ventral dorsal aorta in uninjected controls (n=23 embryos) versus *piezo1;piezo2a* crispants (n=32 embryos), corresponding to the representative images shown in (**b**). Data were collected from two independent experiments and presented as mean ± SD. Statistical significance was determined by two-tailed unpaired *t*-test.

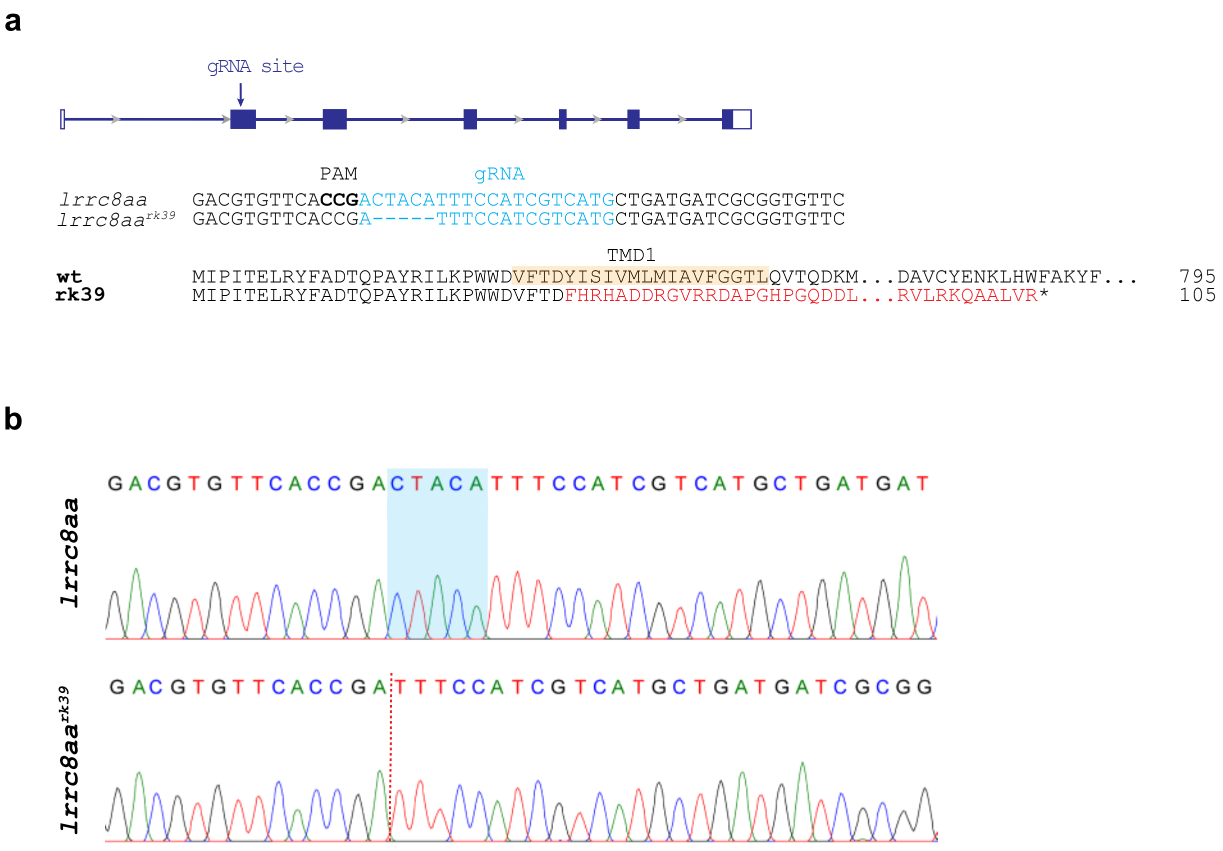


**Supplementary Fig. 18. CRISPR/Cas9-induced mutation in zebrafish *lrrc8aa* gene. a** Zebrafish *lrrc8aa* gene structure, gRNA binding site (in blue), *lrrc8aa$^{rk39}$* allele, Lrrc8aa wild type (795 aa) and Lrrc8aa$^{rk39}$ (truncated at 105 aa) protein structure. The rk39 mutation causes a 5-nt delition which leads to a frameshift after Asp29 and a premature termination codon at amino acid 105 after 76 missense amino acids. TMD1, transmembrane domain. **b** Sequence read showing 5-nt deletion (shadowed area in wild type allele) in lrrc8aark39 allele (red dotted line).
